# CircNFATc3 promotes Fetal Hemoglobin induction by regulating let-7b/GATA2 axis

**DOI:** 10.1101/2025.10.19.682992

**Authors:** Mandrita Mukherjee, Motiur Rahaman, Tuphan Kanti Dolai, Nishant Chakravorty

**Affiliations:** School of Medical Science and Technology, IIT Kharagpur, Kharagpur, Paschim Medinipur, West Bengal –721302, India; Department of Hematology, Nil Ratan Sircar Medical College and Hospital, Kolkata, West Bengal-700014, India

**Keywords:** CircRNA, miRNA, ceRNA network, fetal hemoglobin, erythropoiesis

## Abstract

Reactivation of developmentally suppressed fetal hemoglobin (HbF) holds therapeutic promise. However, the post-transcriptional control of regulators that orchestrate γ-globin (HBG1/2) expression is still elusive. This study aims to elucidate the highly understudied role of circular RNA (circRNA), circNFATc3, as post-transcriptional γ-globin regulator. This study evaluates the circNFATc3/let-7b/GATA2 axis and its relationship with HbF induction through computational and experimental molecular and cellular approaches, including gain and loss-of-function studies. These evaluations have revealed that competitive splicing of circNFATc3, at the expense of its linear transcript, sequesters let-7b miRNA, thereby reactivating GATA2-mediated HbF expression. Moreover, transcriptomic profiling of circNFATc3-overexpressed erythroid cells showed that its role is not only restricted to HbF induction but it may play a broader role in erythropoiesis, adding further complexity to the role of this circular transcript. Therefore, our study demonstrates a pivotal role of circNFATc3 as an HbF inducer and places this circRNA as a promising modifier in erythroid transcription programs, opening avenues for novel therapeutic strategies in β-thalassemia.

## INTRODUCTION

Fetal hemoglobin (HbF; ⍺_2_γ_2_) is a dormant form of hemoglobin in adults with immense therapeutic potential in β-hemoglobinopathies. It is a predominant form of hemoglobin at birth, Fetal hemoglobin is encoded by the nearly-identical genes, HBG1 (^G^γ) and HBG2 (^A^γ), situated in the β-globin locus at chromosome 11[1]. Its expression is suppressed developmentally post-birth by complex long-range interactions between transcription factors and the locus control region (LCR) upstream of the β-globin gene cluster, silencing the embryonic (ε) and fetal genes (γ), and activating the adult β-globin gene[2]. Three quantitative trait loci (QTL) have been accounted for the regulation of HbF expression, namely, BCL11A, intergenic region of HBS1L and MYB, and promoter region of HBG gene, identified by genome-wide association studies. Other developmental factors like LIN28B, IGF2BP2 and let-7 family of microRNAs have also been associated with HBG expression[3]. Natural mutations present in these QTLs (e.g., Hereditary Persistence of Fetal Hemoglobin; HPFH) or pharmacological interventions to reactivate HbF have been reported to reduce clinical severity in β-thalassemia or sickle cell disease[4]. A detailed understanding towards these regulatory mechanisms would provide us with valuable insights in understanding the transcriptional circuitry, thereby promoting HbF reactivation as a therapeutic gain in β-hemoglobinopathies.

As the search for new regulators in HbF expression continues, growing evidence has suggested non-coding RNAs (ncRNAs) as instrumental regulators that provide post-transcriptional control of genes by forming competitive endogenous networks of RNAs (ceRNA network). One such emerging member of the ncRNA landscape is circular RNA (circRNA). Once thought of as “scrambled exons” and a byproduct of spliceosomal anomaly, this biomolecule is currently opening up avenues in cancer research and neurodegenerative diseases. Despite the growing interest, only a few studies have explored their role in HbF regulation [5], [6]. Notably, a recent study identified hsa_circ_0008102 as a prognostic biomarker circulating in the peripheral blood of pediatric β-thalassemia patients[7]. Additionally, two studies have shown that circRNAs can participate in ceRNA networks with miRNAs and mRNAs, for example, hsa_circ_100466-miR-19b-3p-SOX6 and hsa_circ_0005245-hsa-miR-425-3p-GATA2, which have been found to be involved in the regulation of HbF expression in β-thalassemia patients[8], [9]. It is therefore evident from the existing literature that there is a major dearth in understanding the physiological state of HbF expression by circRNA-mediated regulation during erythropoiesis.

Sponge-circRNAs in the cytoplasm act as major regulators of microRNAs (miRNAs) in ceRNA network. While the role of miRNAs in globin gene regulation has been proven extensively making them potential therapeutic candidates to regenerate suppressed HbF in β-hemoglobinopathies, their interactions with circRNAs as globin gene modifiers remain largely unknown[10]. miRNAs are recognized as potent erythroid gene modifiers. Early studies revealed that miR-15a/16-1 and miR-150 target the MYB and act as an indirect HbF inducer[11], [12]. In contrast to this, the precursor form of miR-96 directly binds to γ-globin promoter, leading to polycomb repressor complex-mediated silencing of the gene. Additionally, the miR-144/miR-451 cluster has been demonstrated to regulate the key modulators of fetal hemoglobin including BCL11A, the master regulator of HbF and SOX6[13].

circRNAome profiling of cells undergoing erythropoiesis has provided intriguing observations. These biomolecules are not only cell-specific but also stage-specific across different lineages. Moreover, terminally differentiated erythrocytes have been shown to contain the largest population of circRNAs[14]. This begs the question about its abundance and roles they play in modifying the fate of translational output during differentiation. In this study, we attempted to unravel the regulatory role of circRNAs as miRNA decoys, shedding light on how they orchestrate fetal hemoglobin (HbF) expression during erythroid maturation. To address this, we employed established bioinformatic prediction pipelines for circRNA curation alongside experimental validation approaches. These analyses pointed to circNFATc3 (hsa_circ_0000711) as a potential functional sponge of let-7b miRNA, a known γ-globin suppressor in adult erythroblasts. Moreover, genome-wide transcriptomic profiling suggested that this circRNA may program immature progenitors toward erythropoiesis, highlighting the purposeful role of circRNAs in transcriptional regulation rather than as mere byproducts of spliceosomal activity.

## RESULTS

### Prediction and characterization of circular RNAs in erythroblast cells

To determine the distribution and functionality of circular RNAs that act as modifiers in the globin gene expression during erythropoiesis, a ribo-depleted RNA-Seq dataset of Yu et al. was curated from the GEO database, NCBI. This dataset, GSE121992, consisted of the whole transcriptomic profile of maturing erythroblasts having differential fetal hemoglobin expression (Fig. 1a)[15].

**Figure 1:**
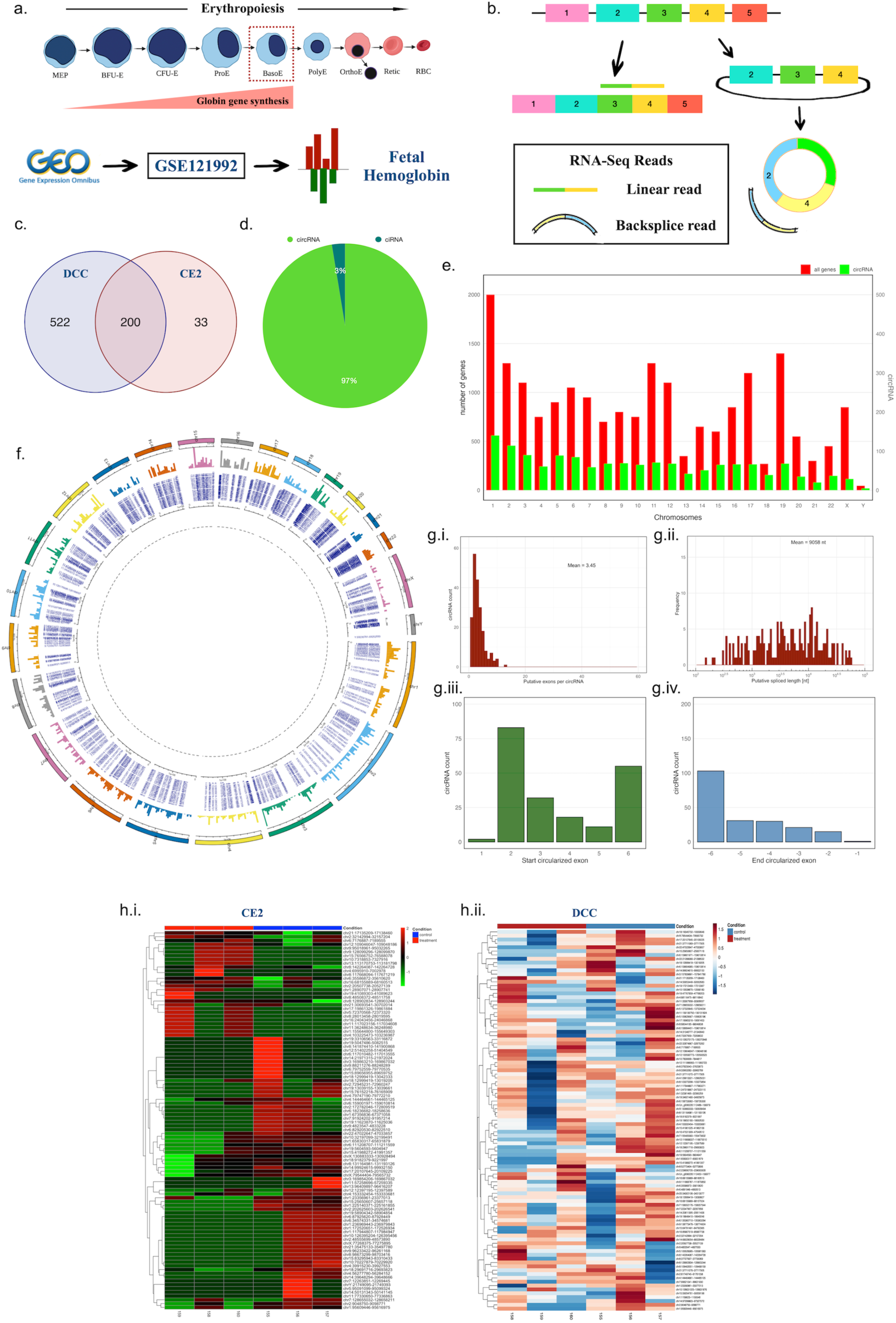
Prediction and characterization of circRNAs in erythroblast cells. **a.** Schematic representation of the total RNA-Seq dataset of basophilic erythroblasts that harbours globin gene expression. **b.** Diagrammatic representation of forward and back splicing recognised by circRNA prediction tools DCC and CE2. **c.** Venn diagram showing the common circRNAs obtained from both tools. **d.** Pie chart showing the distribution of exonic and intronic circRNAs. **e.** Chromosome-wise distribution of circRNAs. **f.** Circos plot showing a detailed view of circRNA distribution and their expression. **g.** Characterizations showing **i.** exon usage **ii.** circRNA length **iii.** start circularisation exon and **iv.** end circularisation exon. **h.** Heatmaps showing expression of the top differentially expressed circRNAs obtained from **i.** CE2 and **ii.** DCC.

Two different tools, Circexplorer2 (CE2) and DCC, were implemented to detect global circRNA expression (Fig. 1b) and their outputs have been provided in Supplementary data 1. To further filter out false positives, a cut-off was applied where circRNAs with at least 2 junction reads in at least 2 biological replicates were considered for further analysis. Initial prediction showed that DCC predicted 49,013 circRNAs and CE2 predicted 6620 circRNAs. After applying the cut-off of at least 2 reads from 2 biological samples, to filter out false positives, 722 and 233 circRNAs remained that were predicted from DCC and CE2, respectively, and there were 200 circRNAs that were overlapping among the two groups (Fig. 1c). Among them, 194 (97%) circRNAs were derived from exon-exon junctions and only 6 (3%) circRNAs were derived from introns (Fig. 1d).

Various characterizations were performed to study circRNA generation and their distribution in erythroid cells. It was observed that circRNAs were generated across all chromosomes (Fig. 1e). To comprehensively understand the distribution of circRNAs across the genome of erythroblasts, the circos plot was adopted for an insightful visualisation. The outer track represents the chromosome numbers, each denoted by a different colour. The immediate inner circle displays the relative expression levels of the circRNAs. Following that, the innermost label track shows the circRNA coordinates with their start and end position. The high density in certain regions of the circle indicates clusters of circRNAs, suggesting hotspots for circRNA biogenesis, for instance in chromosomes 19, 21, and 22 (Fig. 1f).

The circRNAs were further characterised based on their size and exon usage. On average, 3.45 exons were predicted to be used by circRNAs, with a minimum of one and a maximum of 39 exons, as determined by CE2 annotation, based on linear exon usage. The average length of circRNAs was 9058 nucleotides. CircRNAs almost always prefer the second exon to circularize, which is in line with previous findings, where cells of the hematopoietic lineage barely use the first exon for circularisation[14], [16]. It was also observed that exons located further upstream from the last exon were more frequently involved as the end circularized exons. It can be interpreted that there is a strong preference for certain exons or regions of the transcript to be more prone to circularisation, indicating circRNA generation as not a random event (Fig. 1g.i-iv).

CircRNAs that were differentially expressed, annotated from both CE2 and DCC, have been identified, and the significance of circRNAs was considered when their |Log2FoldChange|>1.0 and p-value <0.05. The top differentially expressed genes were plotted in a heatmap (Fig. 1h i-ii). Among the differentially expressed circRNAs derived from both tools, the most upregulated circRNA obtained was hsa_circ_0000711 (chr16: 68155889-68160513), and the most downregulated circRNA was hsa_circ_0005729 (chr18:29691716-29693823).

### Validation of circular RNAs and circRNA-miRNA-mRNA network construction

To validate the predicted circular RNAs, a high fetal hemoglobin expressing K562 cell model was developed using hydroxyurea (HU) drug. Leishman stain showed progressively enlarged cells with reduction in nuclear size (Fig 2a). The flow cytometry study of Erythropoietin (EPO) treated K562 cells showed CD71 and CD235a positive population increasing with time (0h, 48h, 72h) indicating erythroid differentiation (Fig 2bi-ii). Following that, cells were subjected to increasing doses of Hydroxyurea (HU), and it was observed that γ-globin transcription was significantly highest at a dosage of 100μM concentration (Fig 2c). Immunoblot has also shown significant induction of HBG at this concentration (Fig 2d). Next, RT-qPCR was performed to validate the predicted circRNAs in the high HbF K562 cells and it was observed that hsa_circ_0000711 (circNFATc3 derived from *NFATC3* gene in Chromosome 16q22.1) and hsa_circ_0075796 (circDEK derived from *DEK* gene in Chromosome 6p22.3) were significantly upregulated in high HbF condition (Fig. 2e).

**Figure 2:**
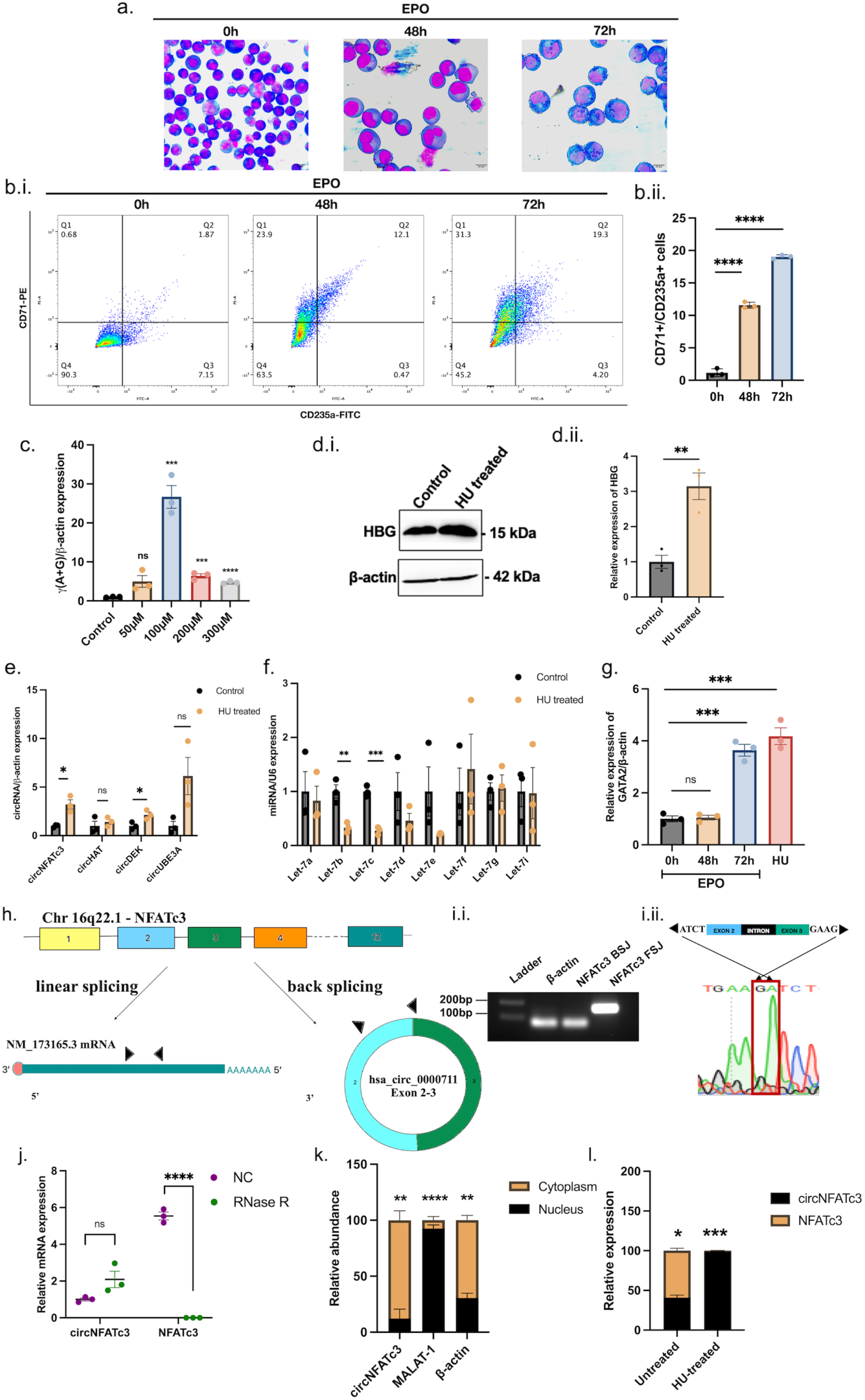
Validation of circRNAs and circRNA-miRNA-mRNA network construction. **a.** Leishman stain showing morphology of EPO-differentiated K562 cells at 0h, 48h and 72h. **b.i.** Flow cytometry plot shows CD71+ and CD235a+ populations at 0h, 48h, 72h to assess erythroid differentiation of EPO-treated K562 cells. **b.ii.** Bar graph representing flow cytometry data of EPO-treated K562 cells undergoing erythroid differentiation. **c.** RT-qPCR analysis of γ-globin mRNA expression at different concentrations of HU-treated K562 cells. **d.i.** Immunoblot showing HBG protein expression in HU-treated and control K562 cells. d.ii. Bar graph showing the quantitative expression of HBG against β-actin. **e.** qRT-PCR analysis of predicted circRNAs in HU-treated cells. **f.** qRT-PCR analysis of let-7 family of miRNAs in high HbF cells. **g.** RT-qPCR of GATA2 in EPO (0h, 48h, and 72h) and HU-treated K562 cells. **h.** Schematic diagram showing the linear and circular isoforms transcribed from NFATc3 gene locus and mapping of the divergent and convergent primers. **i.i.** PCR validation of circNFATc3 using divergent primers. **i.ii.** Sanger sequencing showing the backspliced junction of circNFATc3. **j.** qRT-PCR analysis showing resistance of circNFATc3 against RNase R digestion. **k.** qRT-PCR showing cytoplasmic abundance of circNFATc3. **l.** qRT-PCR showing preferential backsplicing of NFATc3 in HU-treated K562 cells. All experiments were performed in three biological replicates and graphical data are represented as Mean ± SEM, *p≤0.05, ***p≤0.01, ***p≤0.001, ****p<0.0001.

We next examined miRNAs with opposite expression profiles that could potentially interact with the positively regulated circRNAs under high HbF conditions. One of the most notable interactions obtained from circBank was circNFATc3 and let-7b interaction (Supplementary Data 2). Let-7 profiling studies by RT-qPCR was next conducted and let-7b and 7c emerged as the most significantly downregulated members (Fig. 2f). While Let-7a and let-7d has modest downregulation, it was non-significant. Next, targets of let-7b were predicted using MultiMiR and GATA2 emerged as one of the predicted targets of let-7b (Supplementary Data 3). GATA2 mRNA expression was also seen to be gradually increasing in EPO-treated cells at 72h, indicating its involvement in early erythropoiesis, which further increased upon HU treatment, indicating its possible role in γ-globin transcription (Fig. 2g).

Next, the circNFATc3 characteristics were further investigated. This circRNA is known to express from the NFATc3 gene locus that consists of 12 exons on chromosome 16q22.1. The circular variant is transcribed by backsplicing exons 2 and 3. This circRNA is reported in circBase (www.circbase.org), and the size is 1298 bp, and named as hsa_circ_0000711 (circBank ID: hsa_circNFATC3_004) (Fig. 2h)[17]. Next, PCR was performed using divergent and convergent primers against NFATc3 isoforms using cDNA as a template and compared them against β-actin. Results showed divergent primers amplified only the circular isoform, thus detecting and amplifying the backsplice junction, and convergent primers amplified the linear isoform of the NFATc3 gene as expected (Fig. 2i.i). Paired-end Sanger sequencing of the PCR product obtained from the divergent primers confirmed the presence of circNFATc3, by identifying the unique backsplice junction of the circRNA (Fig. 2.i.ii). For specific detection of circNFATc3, RNase R digestion of total RNA was performed. All linear isoforms are expected to be digested, sparing the circular structures which are resistant to exonuclease activity. As expected, a significant decrease in expression of the linear NFATc3 isoform was observed in the RNase R-treated group, while the expression of the circular isoform remained unchanged (Fig. 2j).

The abundance of circNFATc3 in subcellular fractions was assessed. It was found that circNFATc3 was significantly enriched in the cytoplasmic fraction. This indicates its major participation in cytoplasmic transcriptional regulatory network (Fig. 2k). The next assessment entailed the effect of HU treatment on NFATc3 splicing. RT-qPCR analysis demonstrated that in untreated EPO-induced K562 cells canonical splicing prevailed, but when treated with 100μM HU backsplicing induced circNFATc3 was significantly upregulated at the expense of its linear isoform whose mRNA expression decreased significantly (Fig. 2l). Taken together, these experiments confirm the circular structure of circNFATc3.

### cirNFATc3-let-7b-GATA2-γ-globin axis in primary erythroid cells

To study the stage-specific function of circNFATc3 and its modulatory effect on let-7b, GATA2 and HBG, CD34+ HSPCs were treated with 100µM Hydroxyurea containing erythroid differentiation media for 8 days. Cells were harvested at two time points, Day 4 and Day 8. From our *in-silico* analyzes, it was observed that circNFATc3 was significantly expressed in the basophilic erythroblast phase which was achieved at 8th day of differentiation. FACS analysis on these two time points have shown significant increase in double positive CD71+/CD235a+ population, indicating active erythropoiesis (Fig 3a.i-ii). Cells were subjected to Leishman stain to study their morphology and it was observed that at day 4 the cells were typically smaller in size. By 8^th^ day, cells had increased by size, with shrunken nuclei, resembling basophilic erythroblasts (Fig. 3b). RT-qPCR studies showed significant increase in expression of circNFATc3, GATA2 and HbF by Day 8 while let-7b was significantly downregulated (Fig 3c. i-iv). It was also concluded that circNFATc3 expression was not cell line specific but had a significant expression in high HbF expressing primary erythroid cells as well. Therefore, we selected the circNFATc3-let-7b-5p-GATA2 axis to further unfold its role in HbF regulation.

**Figure 3.**
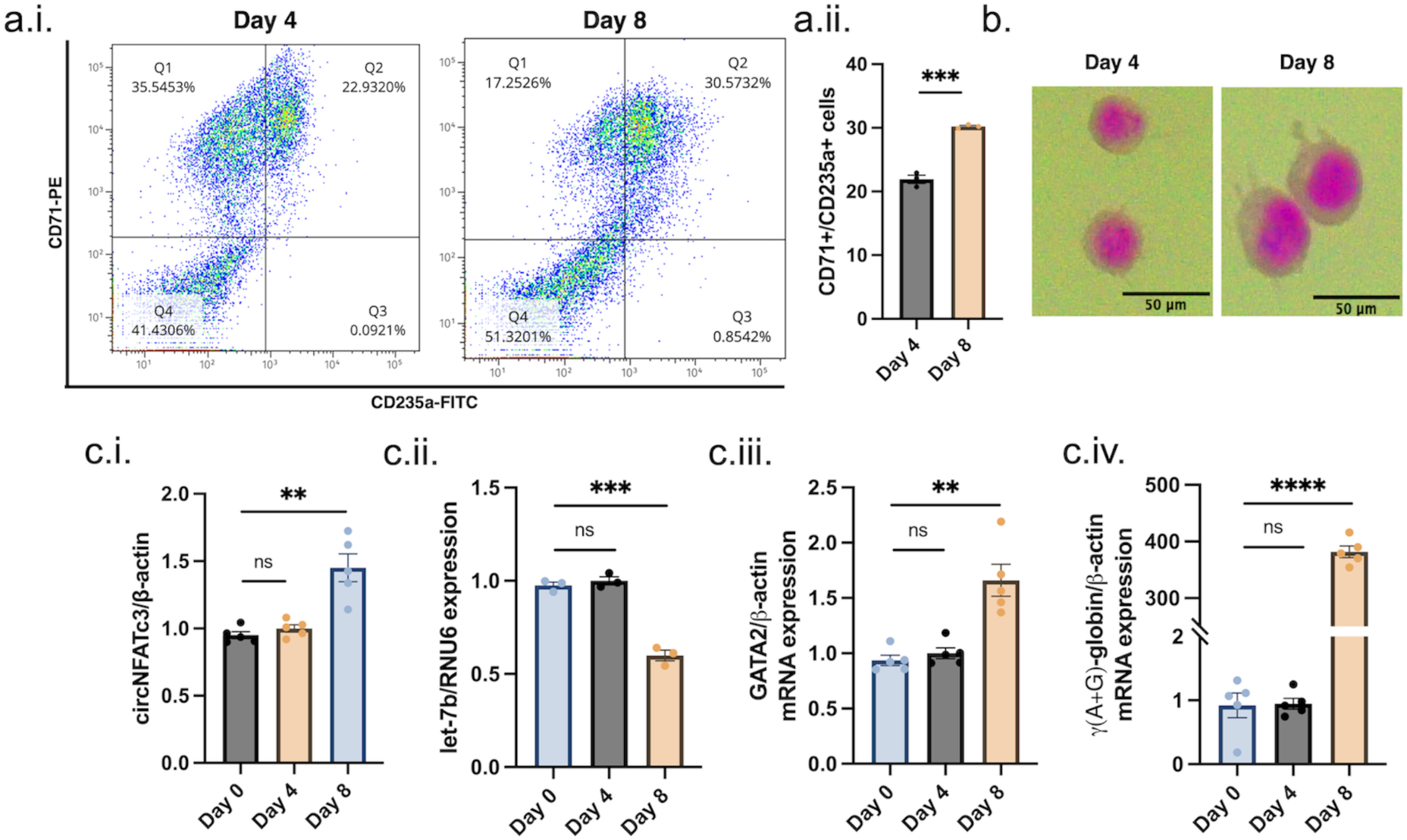
ceRNA axis in primary erythroid cells. **a.i.** Flow cytometry analysis of differentiation markers CD71 and CD235a double positive population at different time points. **a.ii.** Bar graph of representative flow cytometry data showing erythroid differentiation of CD34+ HSPCs at different time points. **b.** Leishman stain showing the morphological differences between ex vivo differentiating primary erythroid cells at Day 4 and Day 8. **c.** RT-qPCR analysis of **i.** circNFATc3, **ii.** Let-7b, **iii.** GATA2 and **iv.** γ(A+G)-globin. All experiments were performed in three biological replicates and graphical data are represented as Mean ± SEM, *p≤0.05, ***p≤0.01, ***p≤0.001, ****p<0.0001.

### hsa-let-7b-5p represses gamma globin expression by GATA2-mediated silencing

To explore the ability of let-7b as a fetal hemoglobin repressor, miRNA mimics were ectopically overexpressed in K562 cells and further analysis was performed (Fig 4a). Cell viability assay showed no detrimental effect on cell growth upon transfection of mimics and mimic negative control up to 96 hours (Fig 4b). Next, qRT-PCR assessment showed significant overexpression of let-7b mimics post-transfection. Furthermore, we also observed that GATA2 and γ-globin mRNA transcription was significantly reduced, while, circNFATc3 and β-globin expression remained unchanged (Fig 4c.i-v). Immunoblot studies showed similar findings where GATA2 and HBG protein expression was significantly suppressed (Fig 4d.i-ii).

**Figure 4:**
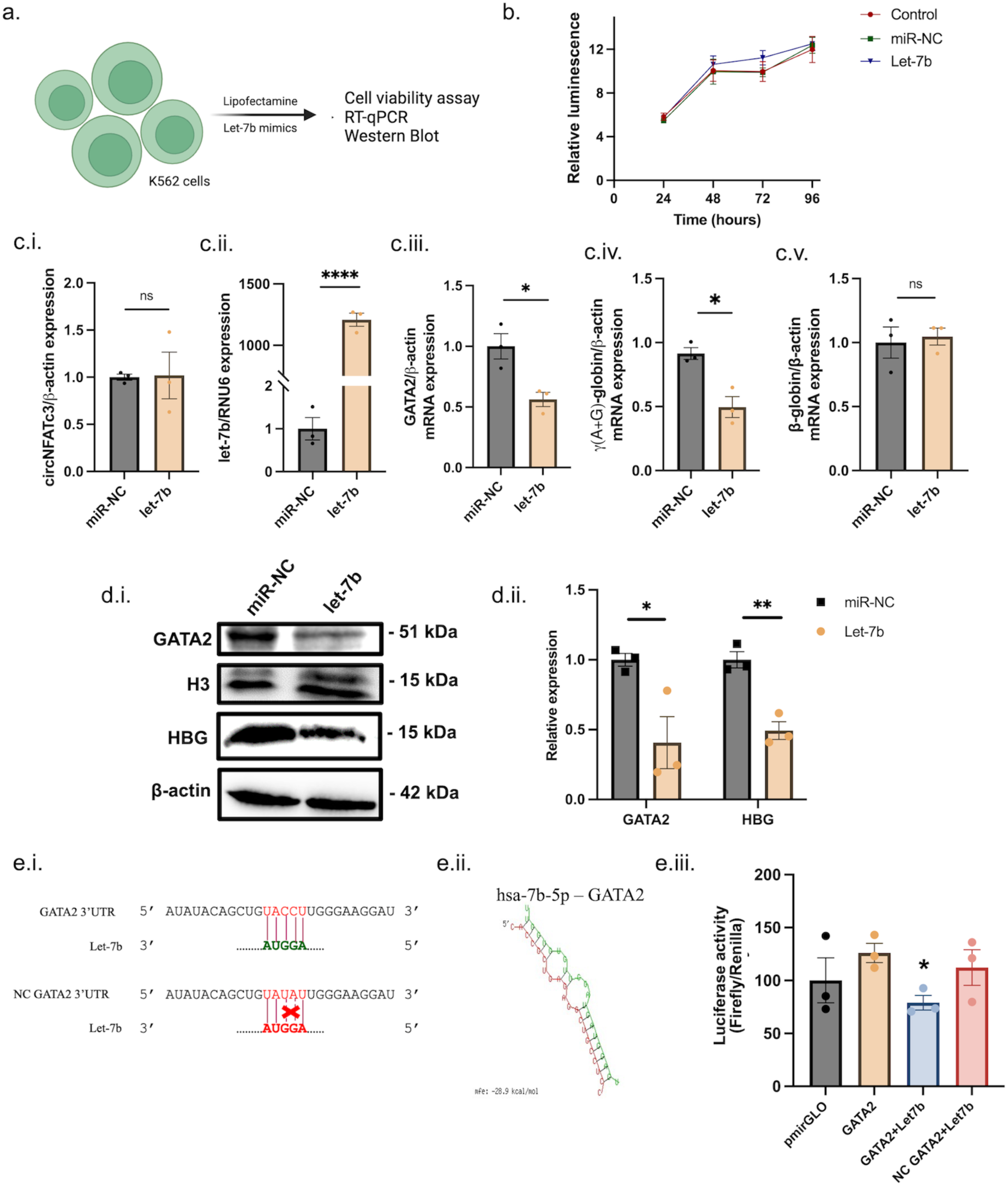
Effect of Let-7b overexpression on fetal hemoglobin expression. **a.** Schematic diagram showing transfection of miRNA mimics (let-7b mimic and miR-NC) in K562 cell line. **b.** Cell Titer-Glo assay results showing proliferation of the cells at different time points post miRNA transfection. **c.** qRT-PCR analysis of **i.** Let-7b, **ii.** circNFATc3, **iii.** GATA2, **iv.** γ(A+G), and **v.** β-globin. **d.i.** Immunoblot analysis showing expression of GATA2, Histone H3, HBG, and β-actin in miRNA transfected K562 cells, **d.ii.** Relative expression analysis of immunoblot bands of GATA2 and HBG. **e.i.** Schematic showing GATA2 3’UTR sequence seed matching with let-7b and mutated GATA2 seed region mismatched with let-7b, **ii.** Schematic showing RNAhybrid output of hsa-let-7b-5p and GATA2 interaction (MFE=-28.9kcal/mol) **iii.** Relative luciferase activity in HEK293T cells co-transfected with let-7b mimics and luciferase reporter vector. All experiments were performed in three biological replicates and graphical data are represented as Mean ± SEM, *p≤0.05, ***p≤0.01, ***p≤0.001, ****p<0.0001.

Furthermore, to assess if let-7b directly targeted GATA2, we predicted the binding site of the miRNA in the GATA2 3’UTR region by RNAhybrid (Fig 4e.i-ii). Next, we performed a luciferase reporter assay to investigate the interaction of Let-7b and GATA2. Our results demonstrated that let-7b overexpression significantly repressed the firefly luciferase activity when the 3’UTR sequence of GATA2 was cloned, while no repression was exhibited when the mutant 3’UTR was cloned (Fig 4e. iii). Together, our results indicate that let-7b miRNA directly targets and silences GATA2, and subsequently represses fetal hemoglobin expression. The unchanged expression of circNFATc3 upon let-7b overexpression warranted further investigation. Taken together, these findings suggest let-7b miRNA to be a repressor of GATA2 and HbF.

### Overexpression of circNFATc3 reactivates fetal hemoglobin

To establish the effect of circNFATc3 on fetal hemoglobin regulation, circNFATc3 overexpression was performed *in vitro.* pcDNA3.1(+) circRNA mini vector was used for overexpressing while the linear transcript of the circRNA sequence served as our negative control (Fig. 5a). Following transfection, the viability of cells was monitored and observed that there was no significant difference in growth (Fig. 5b).

**Figure 5:**
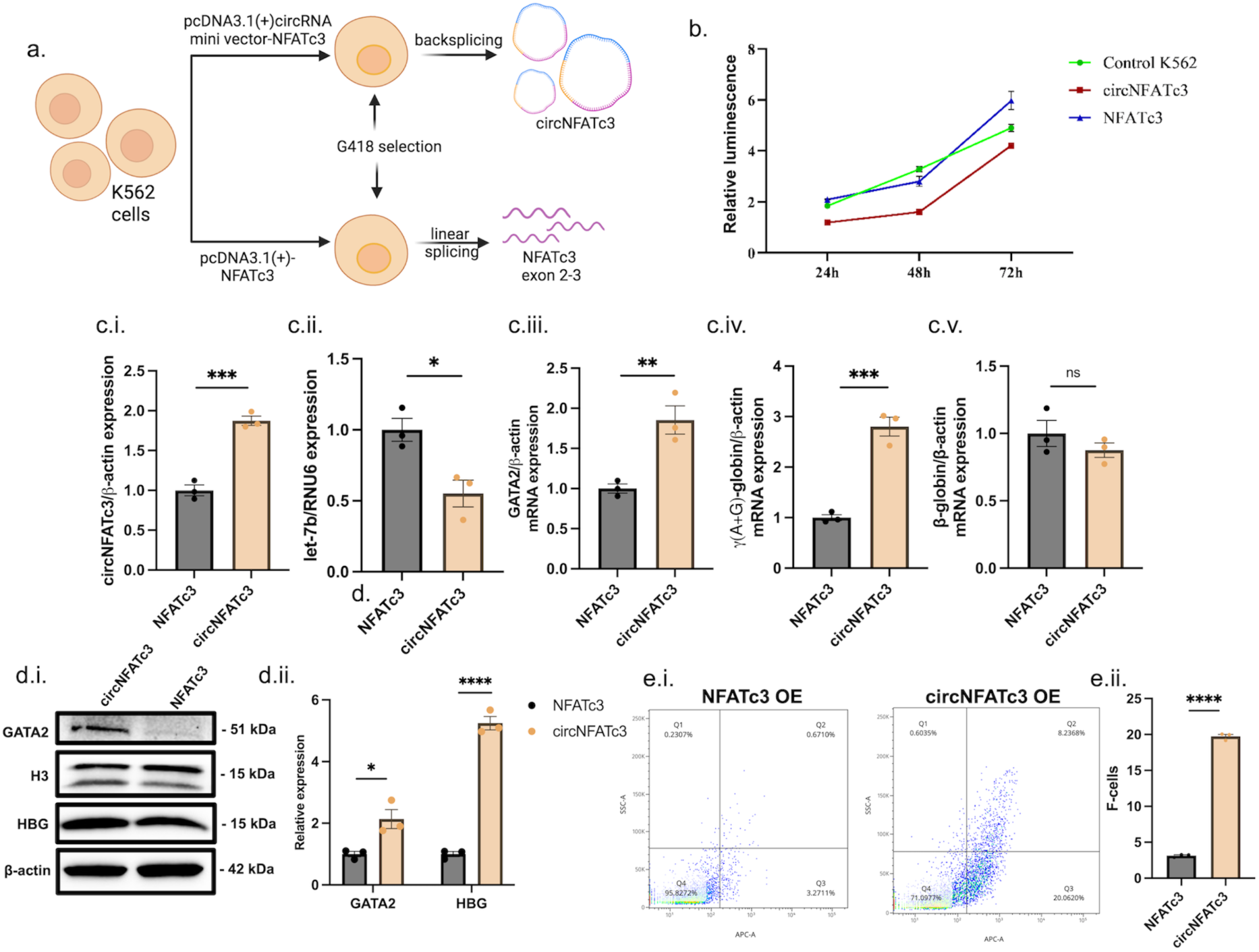
Overexpression of circNFATc3 induces fetal hemoglobin expression. **a.** Schematic diagram showing K562 cells stably overexpressed with circNFATc3 and NFATc3 exon 2-3. **b.** Cell Titer-Glo assay results showing proliferation of the cells at different time points post-transfection of pcDNA3.1(+)-circRNA mini vector-NFATc3 exon 2-3 and pcDNA3.1(+)-NFATc3 exon 2-3 **c.** qRT-PCR analysis of **i.** circNFATc3, **ii.** let-7b, **iii.** GATA2, **iv.** γ-globin, **v.** β-globin. **d.i.** Immunoblot analysis showing expression of GATA2, H3, HBG, β-actin. **ii.** Relative expression analysis of immunoblot bands of GATA2 and HBG. **e.i.** Flow cytometry analysis of F-cells overexpressing circNFATc3 and NFATc3 exon 2-3. **ii.** Bar graph of representative flow cytometry data showing percentage F-cell population. All experiments were performed in three biological replicates and graphical data are represented as Mean ± SEM, *p≤0.05, ***p≤0.01, ***p≤0.001, ****p<0.0001.

Next, we investigated the expression profiles of the circRNA/miRNA/mRNA axis upon circNFATc3 overexpression. Upon significant upregulation of circNFATc3, let-7b was significantly downregulated, while GATA2 and γ-globin were significantly upregulated. β-globin mRNA expression remained unchanged, confirming an independent regulatory effect of circNFATc3 on γ-globin regulation (Fig 5c. i-v). Immunoblot studies showed significant upregulation of GATA2 and HBG expressions (Fig 5d. i-ii). The F-cell population in the circNFATc3 overexpressed cells also showed a significant increase compared to the linear transcript overexpressing cells (Fig. 6e. i-ii). Taken together, these findings show circNFATc3 as an inducer of HbF.

**Figure 6:**
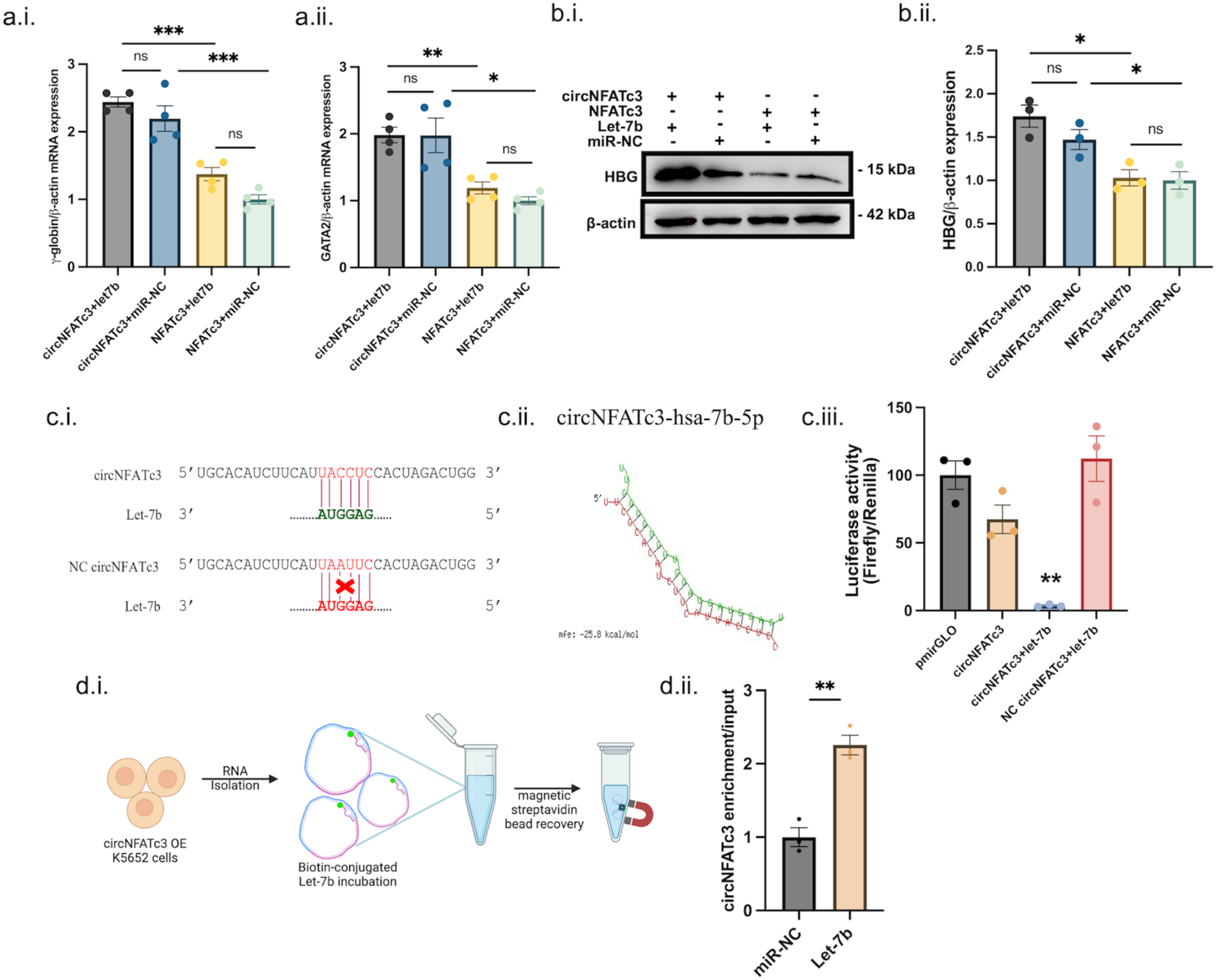
circNFATc3 sponges Let-7b and induces fetal hemoglobin. **a.** qRT-PCR analysis of **i.** GATA2 and **ii.** HBG in let-7b and miR-NC transfected circNFATc3 and NFATc3 overexpressing cells. **b.i.** Immunoblot analysis of HBG in overexpressed circNFATc3 and NFATc3 cells co-transfected with let-7b mimics and miR-NC. **ii.** Bar graph showing relative expression analysis of immunoblot bands of HBG. **c.i.** Schematic showing circNFATc3 sequence seed matching with let-7b against and mutated circNFATc3 seed region mismatched with let-7b **ii.** Schematic showing RNAhybrid output of circNFATc3 and hsa-let-7b-5p interaction (MFE=-25.8kcal/mol). **iii.** Relative luciferase activity in HEK293T cells co-transfected with let-7b mimics and luciferase reporter vector. **d.i.** Schematic showing let-7b-mediated RNA-pull down assay methodology. **ii.** RT-qPCR showing enriched circNFATc3 expression. All experiments were performed in three biological replicates and graphical data are represented as Mean ± SEM, *p≤0.05, ***p≤0.01, ***p≤0.001, ****p<0.0001.

### circNFATc3 sponges hsa-let-7b-5p to induce fetal hemoglobin

circNFATc3 is primarily expressed in the cytoplasm; therefore, we speculated that it may act as a miRNA sponge. To further investigate this, let-7b mimics and negative control were transfected in overexpressing circNFATc3 and linear NFATc3 cells. It was observed that the transfection of let-7b mimics or miR-NC had no effect on both GATA2 and γ-globin transcription, and they were significantly upregulated in circNFATc3 overexpressing cells (Fig 6a. i-ii). Similar observations were seen in immunoblot analysis of HBG (fig 6b. i-ii). Linear NFATc3 and let-7b or miR-NC also showed similar non-significant change in expression, although the expression of HBG and GATA2 were significantly higher in circNFATc3 overexpressing cells co-transfected with let-7b or miR-NC.

There was no significant difference in expression of HBG and GATA2 when let-7b and miR-NC were transfected in NFATc3 overexpressing cells. To explore circNFATc3 sponge activity, the miRNA response element (MRE) for hsa-let-7b-5p was identified in the circRNA from circBank and RNAhybrid, and luciferase reporter constructs were co-transfected with miRNA mimics in HEK293T cells. It was observed that luminescence activity of firefly luciferase was significantly reduced in pmirGLO vector cloned with circNFATc3 seed sequence of let-7b, compared to vector cloned with mutant seed region (Fig 6c. i-iii). To further prove the direct interactions between circNFATc3 and let-7b, a biotin-streptavidin-based RNA-pull down assay was conducted. RT-qPCR analysis revealed circNFATc3 to be significantly enriched compared to miRNA negative control (Fig 6e. i-ii). Taken together, these results clearly indicate circNFATc3 functions as a GATA2-mediated fetal hemoglobin inducer by acting as a sponge for hsa-let-7b-5p miRNA.

### Knockdown of circNFATc3 suppresses fetal hemoglobin

To establish the effect of circNFATc3 silencing on HbF, as shown in the schematic, siRNA specific to the backsplice junction of circNFATc3 was custom-designed and the dosage was optimised (Fig 7a). At first, varied concentrations of siRNA were transfected *in vitro* and monitored for 72 hours, and it was seen that there were no significant changes in cell growth compared to control (Fig 7b). Following that, RT-qPCR was performed to study the dose-dependent effect of siRNA on circNFATc3 and γ-globin transcription (Fig 7c. i-ii). It was seen that 50 pmol siRNA showed maximum silencing after which increasing concentration showed minimized silencing, possibly due to reduced silencing efficiency[18]. GATA2 mRNA levels were shown to be significantly suppressed at 50 and 100 pmol concentration of siRNA (Fig 7c.iii). Following that, immunoblot assesment of HBG demonstrated a similar expression pattern as RT-qPCR where 50 pmol siRNA showed better silencing than 100 pmol concentration (Fig 7d. i-ii). Taken together, the silencing studies clearly revealed the influence of circNFATc3 levels on HbF expression.

**Figure 7:**
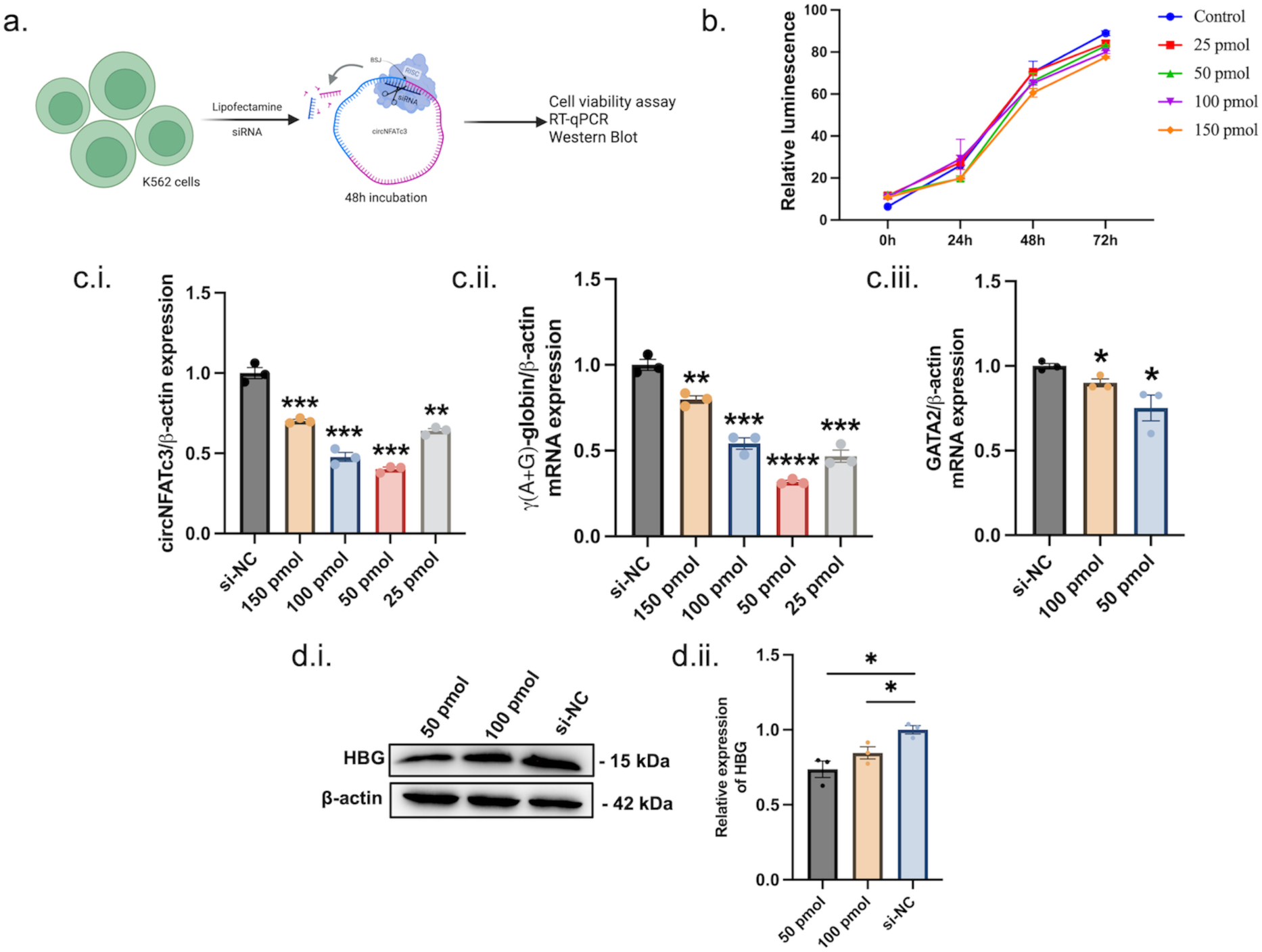
Knockdown of circNFATc3 silences fetal hemoglobin. **a**. Schematic diagram showing transfection of siRNA (si-circNFATc3 and si-NC) in K562 cell line. **b.** Cell Titer-Glo assay results showing proliferation of the cells at different time points post-transfection of different concentrations of siRNA. **c.** RT-qPCR analysis shows **i.** dosage optimisation of siRNA on circNFATc3, **ii.** γ-globin and **iii.** GATA2. **d.i.** Immunoblot analysis of HBG expression upon siRNA-mediated circNFATc3 silencing and **ii.** Relative expression analysis of immunoblot bands of HBG in si-circNFATc3 and si-NC transfected cells. All experiments were performed in three biological replicates and graphical data are represented as Mean ± SEM, *p≤0.05, ***p≤0.01, ***p≤0.001, ****p<0.0001.

### circNFATc3 influences broader erythroid programs and fetal hemoglobin regulation

Morphology analysis by Leishman staining exhibited cells, overexpressing circNFATc3, were undergoing active erythropoiesis, where basophilic erythroblasts were largely populated (45%) followed by polychromatic erythroblasts (29%). Some cells had condensed nuclei presenting basophilic, polychromatic and orthochromatic stages. The polychromatic stage shows condensed and fragmented nuclei which was more profound in the orthochromatic stage where cell size had also greatly reduced. Their rate of proliferation was significantly high compared to the NFATc3 exon 2-3 linear transcript, which resembled a closer morphology to untreated K562 cells. (Fig. 8.a i-iii). Expression profile of key erythroid genes ALAS2, GATA1 and BAND3 were significantly upregulated in circNFATc3 overexpressing cells (Fig 8b. i-iii)

**Figure 8:**
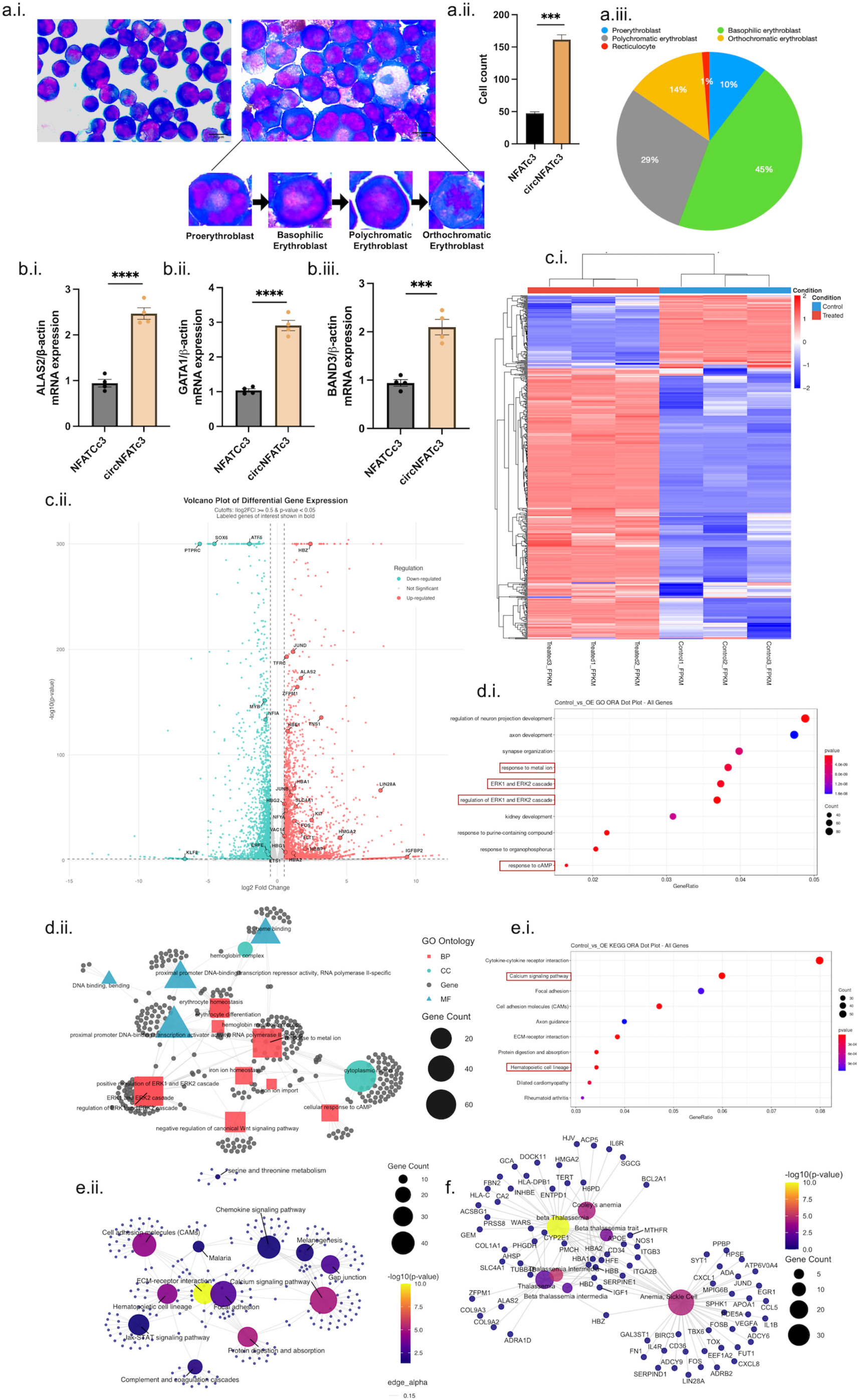
circNFATc3 influences broader erythroid programs and fetal hemoglobin regulation. **a.i.** Leishman stain showing the morphological differences between circNFATc3 overexpressing cells and its linear transcript. **ii** Bar graph showing cell count of circNFATc3 and NFATc3 overexpressed cells **iii.** Pie chart showing the distribution of different erythropoietic stages. **b.** RT-qPCR studies showing key erythroid genes expression of **i.** ALAS2, **ii.** GATA1 and **iii.** BAND3. **c.i.** Heatmap showing differentially expressed genes upon circNFATc3 overexpression. **ii.** Volcano plot showing significant differentially expressed genes (|log2FC|=0.5, p-value<0.05). **d.i.** Gene Ontology ORA Dot Plots for Significant DGE Protein Coding Genes. This plot displays the top 10 enrichments for all the significant DGE protein coding genes (up and down regulated together) group wise. The results are displayed taking gene ratio on the x-axis and GO term on the Y axis. Size of the dot represents the numbers of genes in the enrichment pool for each GO term. Colour of the dot represents the p-value from enrichment result. (Red: Min and Blue: Max). **d.ii.** Gene Ontology ORA Gene-Concept Network Plot for Significant DGE Protein Coding Genes. This plot depicts the linkages of all the significant protein coding DGE genes (up and down-Regulated) and biological concepts (e.g. GO terms) as a network for each group. **e.i.** KEGG Pathway ORA Dot Plot for Significant DGE Protein Coding Genes. This plot displays the top 10 enrichments for all the significant DGE protein coding genes (up and down regulated together) group wise. The results are displayed taking gene ratio on the x-axis and KEGG term on the Y axis. Size of the dot represents the numbers of genes in the enrichment pool for each KEGG term. Colour of the dot represents the p-value from enrichment result. (Red: Min and Blue: Max). **e.ii.** KEGG Pathway ORA Gene-Concept Network Plots for Significant DGE Protein Coding Genes. This plot depicts the linkages of all the significant protein coding DGE genes (up and down-Regulated) and biological concepts (e.g. KEGG terms) as a network for each group. **f.** Disease Ontology ORA Gene-Concept Network Plots for Significant DGE Protein Coding Genes. This plot depicts the linkages of all the significant protein coding DGE genes (up and down-Regulated) and biological concepts (e.g. DO terms) as a network for each group. All experiments were performed in three biological replicates and graphical data are represented as Mean ± SEM, *p≤0.05, ***p≤0.01, ***p≤0.001, ****p<0.0001.

Gene profiling of K562 erythroid cells was performed from 3 treated (circNFATc3 overexpressed) and 3 control (linear NFATc3 overexpressed) samples, and we identified 5508 significant differentially expressed genes (Supplementary Data 4). Heatmap analysis exhibited distinct gene expression patterns (Fig 8c. i). A more detailed visualisation in the volcano plot revealed some key genes known to be involved in HbF regulation and erythroid differentiation (|log2FC|>0.5 and p-value<0.05) (Fig 8c. ii).

Gene ontology (GO) provided some fundamental insights regarding the biological processes (BP), cellular component (CC) and molecular functions (MF) and their associated genes significantly enriched in HbF synthesis and erythroid maturation (Fig 8d. i-ii and Supplementary data 5). KEGG pathway ORA provided a comprehensive understanding of significantly enriched pathways involved in erythropoiesis and HbF expression (Fig 8e. i-ii and Supplementary data 6). DisGenNet analysis has been conducted to study the gene-disease association and 1054 diseases were significantly enriched (p<0.05). The disease ontology network has been plotted focusing on β-hemoglobinopathies to achieve an understanding of the dysregulation of globin genes and associated modulators of erythropoiesis (Fig 8f and Supplementary data 7).

## DISCUSSION

Expression of human fetal hemoglobin (HbF) is suppressed after birth, and its regeneration in adults holds promise in reversing the severity of the disease in patients with β-hemoglobinopathies[19]. The fetal to adult switch in hemoglobin, post-birth, is a complex and tightly regulated mechanism involving numerous transcription factors and long-range chromatin loop interactions[20]. As the search for new modulators of fetal hemoglobin continues, the family of non-coding RNAs (miRNAs, circRNAs and lncRNAs) have emerged as a potent regulator of this switching mechanism. The cytoplasm of erythroid cells harbours one of the most complex and intricate competitive endogenous networks of RNAs (ceRNAs) that play a vital role in influencing the translation output of genes[21]. Among them, circular RNAs have gained attention as a class of non-coding RNAs for their unique circular structures that have better resistance to exonuclease, therefore making them potential candidates for RNA therapeutic interventions like stable protein translation, vaccine formulations, and miRNA or protein sponges[22].

To this end, various microRNAs have been identified that are directly or indirectly linked to hemoglobin switching mechanism[23]. One of the most crucial family of miRNAs, the let-7 family act as potent suppressors of fetal hemoglobin in adult erythroblasts and are downregulated in HPFH[24]. Let-7a and let-7b are the most predominantly expressing members, which are developmentally suppressed by LIN28B in fetal erythroblasts. In adults, as the expression of LIN28B declines, let-7 effectively decoys HIC2, thus allowing BCL11A-mediated HbF suppression[25]. This recent study has revolutionized our understanding of the complex regulation of let-7 miRNA and its influence on fetal hemoglobin expression, however, circRNA-mediated let-7 regulation remains largely unknown. To the best of our knowledge, this is the first study to identify let-7 regulation by sponge-circRNAs. Thus, we have unraveled circNFATc3 as a novel regulator of fetal hemoglobin through its role as a ceRNA that decoys let-7b miRNA thereby promoting GATA2-mediated HbF reactivation.

The identification of circNFATc3 through circRNA prediction pipelines (Circexplorer2 and DCC) and its subsequent validation demonstrates a significant advancement in understanding circRNA-mediated regulation of fetal hemoglobin. Our transcriptomic analysis has revealed the abundant presence of circRNAs across chromosomes in erythroid cells, with non-uniform clustered hotspots for circRNA biogenesis, indicating dedicated islands for circRNA generation with definitive regulatory roles. A majority of circRNAs were shown to derive from exonic regions, consistent with established patterns of circRNA biogenesis in erythroid lineage[14]. This may point to a possible maintenance of the proteome after enucleation of erythroblasts, provided an intact internal ribosome entry site (IRES) is present in these circRNAs[16]. Another notable finding is that the circRNAs mostly favors the second exon for circularization, while there no clear preference of the last exon usage, which in concert with previous findings in cells of hematopoietic lineage[14], [16]. Although, no clear explanation has been provided for this preference, the second exon offers both an upstream 3’ acceptor splice site and downstream 5’ donor splice site in the intronic sequences for successful backsplicing, thus providing a positional advantage for the spliceosomal machinery[26]. However, this explains eligibility rather than preference of the second exon.

*In silico* and RT-qPCR validation experiments revealed circNFATc3 (hsa_circ_0000711) as an upregulated circRNA in high fetal hemoglobin condition. It exhibits RNase R resistance due to its circular structure, and its unique backsplice junction was confirmed by Sanger sequencing. The cytoplasmic abundance of circNFATc3 confirms its participation in competitive endogenous RNA networks, where it can effectively sequester miRNAs and modulate the fate of gene expression post-transcriptionally. The competitive splicing mechanism that chooses circular transcript over the linear counterpart of NFATc3 gene, in high HbF conditions, exhibits a sophisticated spliceosomal regulation in erythroid cell.

To further substantiate the presence of circNFATc3/let-7b/GATA2 axis, that have been predicted *in-silico*, RT-qPCR analysis of this axis was performed in primary maturing erythroblast cells expressing high fetal hemoglobin at different stages of differentiation. Intriguingly, circNFATc3 is widely expressed in primary erythroblasts and it upregulates as erythroblasts, expressing high HbF, matured and reached basophilic erythroblast stage on Day 8, which concerted with our *in-silico* discovery, indicating the stage-specific nature of circRNAs[14].

Let-7b overexpression studies revealed significant downregulation of GATA2 and HbF, while HbA (adult hemoglobin; ⍺_2_β_2_) remained unchanged. This supports previously established studies that let-7 mediates γ-globin suppression through distinct regulatory pathways[27]. Furthermore, luciferase reporter assay revealed that let-7b directly interacts with GATA2, a finding that had not been previously reported. Being understudied for its role in HbF induction, GATA2 is an essential part of the “GATA switching factor” along with GATA1 that orchestrates erythropoiesis during development. One study shows that GATA2 facilitates HbF expression by partnering with NF-Y in fetal erythroblasts by binding to the CCAAT box upstream of γ-globin promoter, which eventually gets displaced in adult erythroblasts by BCL11A and GATA1 along with the NuRD complex that supresses HbF post-birth[28]. Furthermore, circNFATc3 expression remained insignificant, thus indicating its upper hand in let-7b regulation by sponging mechanism.

Acting as a putative sponge-circRNA for let-7b in modulating HbF expression, circNFATc3 overexpression studies were carried out for further investigation. This circRNA’s role in hematopoietic lineage has not been unraveled yet, therefore, the emergence of circNFATc3 as a top candidate in high HbF conditions triggered our interest. The overexpression of circNFATc3 led to significant reduction in let-7b expression, while GATA2 and HBG significantly increased, but HbA remained unchanged. These findings indicate circNFATc3 as an independent γ-globin inducer through let-7b sponging, further revealed through its enrichment by bio-let-7b-5p-mediated pull-down and luciferase reporter assays. The HbF inducing properties of circNFATc3 was confirmed when FACS analysis showed significant increase of F-cell population in circNFATc3-overexpressing erythroid cells. Upon co-expression of circNFATc3 and let-7b, HBG and GATA2 were significantly upregulated, which substantiated our hypothesis of circNFATc3 as a decoy of let-7b. One observation that intrigued us was the non-significant effect of let-7b and miR-NC in linear NFATc3 overexpressing cells, considering NFATc3 linear species should not have any influence on let-7b. A possible explanation may be that the linear transcripts can circularise if they contain structural motifs (e.g., complementary introns or RBP-binding sites) that promote backsplicing, and overexpression may amplify this process. However, overexpression systems are prone to generating heterogeneous RNA species, therefore require rigorous validation to confirm circRNA-specific effects[29], [30].

One of the most intriguing findings of the study was the dose-dependent relationship observed in the siRNA-mediated silencing studies of circNFATc3. Optimal silencing of GATA2 and HBG was achieved at 50 pmol concentration of si-circNFATc3 dosage, and a gradual decrease in the silencing efficiency at higher doses of siRNA was observed. Therefore, the principles of ceRNA hypothesis are substantiated where ceRNA binding is governed by stoichiometric ratios rather than on/off switches[31]. The observation that γ-globin expression showed a parallel dose-response relationship to circNFATc3 silencing demonstrates a direct link to the population of ceRNAs, and thus establishes a quantitative aspect to the understanding of these regulatory networks. Although significant, subtle differences in GATA2 expression may have a substantial biological impact on HBG expression.

Morphological analysis revealed distinct phenotypic difference in circNFATc3 overexpressing cells, where they spontaneously matured into erythroblasts without the usage of erythroid-inducing growth factors like EPO. Cells were undergoing active erythropoiesis, where majority were in the basophilic (45%) and polychromatic erythroblast (29%) stages, which strengthens our previous finding, that circNFATc3 expression is pronounced in basophilic stage of primary erythroblasts. Expression profile of key erythroid differentiation factors (GATA1, ALAS2 and BAND3) also exhibited significant upregulation.

While this study contributes to the understanding of circRNAs in HbF regulation, it provides us with a single regulatory axis among many other players that participate in the ceRNA network. For a broader perspective of circNFATc3’s role in HbF regulation and erythroid maturation, genome-wide expression profiling studies have been performed on overexpressed circNFATc3 erythroid cells. Certain HbF inducers were significantly upregulated such as LIN28A[32], NFYA[33], NFIA[34], [35], FLT1[35], etc. while the suppressors like SOX6[36], MYB[37], KLF8[38], etc. were significantly downregulated. The expression profile of circNFATc3 may mirror certain aspects of HPFH that are demonstrated in the Supplementary data 9 and 10. While significant upregulation was observed in iron uptake and erythroid differentiation genes [39], [40], Gene Ontology (GO) studies have shown enrichment of pivotal pathways crucial for erythropoiesis, HbF induction, heme biosynthesis, iron uptake, hemoglobin synthesis, etc. Disease enrichment have also shown associations with β-hemoglobinopathies.

This complex transcriptional reprogramming observed, positions circNFATc3 as a promising modifier of fetal hemoglobin and erythropoiesis. Our findings suggest that circNFATc3 exerts a competitive binding on hsa-let-7b-5p and thereby modulates the expression of GATA2 (Fig 9). As a result, the suppressive activity of let-7b on GATA2 is disrupted, which allows HbF induction, thus ultimately opening avenues to provide therapeutic interventions in β-hemoglobinopathies.

**Figure 9:**
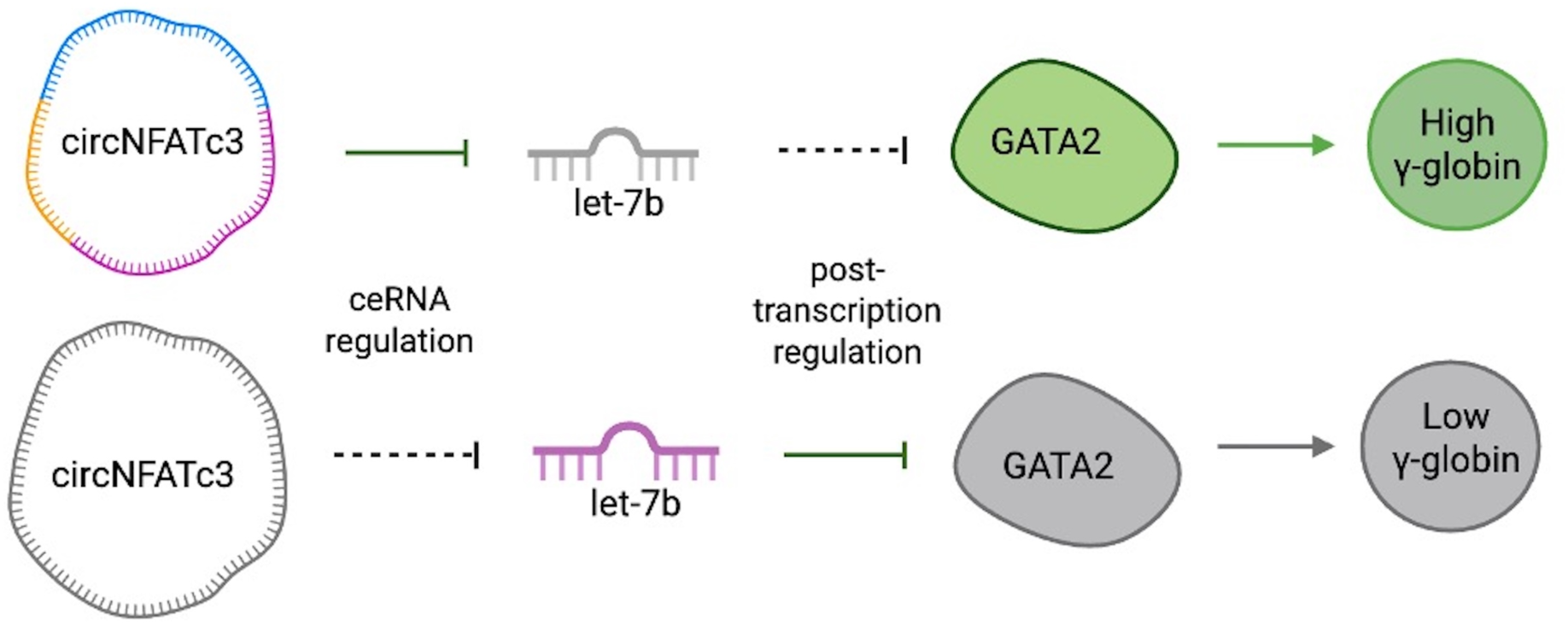
Schematic representation of HbF regulation by circNFATc3/let-7b/GATA2 axis. circNFATc3 overexpression sequesters let-7b, thereby allowing GATA2-mediated HbF induction. In contrast, circNFATc3 silencing allows let-7b-mediated sequestration of GATA2, subsequently repressing HbF.

## MATERIALS AND METHODS

### Bioinformatic prediction of circRNAs and miRNAs

Gene Expression Omnibus (GEO) datasets related to fetal hemoglobin were searched from NCBI (www.ncbi.nlm.nih.gov/geo/). A high-throughput ribosomal RNA-depleted paired-end RNA sequencing GEO dataset of early basophilic erythroblast cells (GSE121992) was selected, which had long reads of 125 nt. The dataset comprised of three low and three high fetal hemoglobin samples, generated by CRISPR/Cas9-mediated knockout of MBD2, a component of the NuRD complex and a repressor of HbF[15]. The raw fastq files of SRR8137155, SRR8137156, SRR8137157, SRR8137158, SRR8137159, and SRR8137160 were retrieved from the SRA repository and aligned to the human reference genome GRCh37/hg19 using STAR version 2.7.11b to detect chimeric reads[41]. Following that, the Chimeric.out.junction files were used for detecting, filtering, and annotating circRNAs using Circexplorer2 (CE2) v2.01 and DCC v0.4.4. GRCh37/hg19 genome annotations were used for both CE2 and DCC[42], [43]. The output.txt file provided us with the chromosome coordinates, exon counts, read counts, etc. Next, circRNA analysis was performed using the DCC tool with the same Chimeric.out.junction file, where the output file was obtained with similar information as Circexplorer2. A cut-off was applied where circRNAs were considered when at least two read numbers were present in at least two biological samples[44]. For differential expression analysis of the high-confidence circRNAs, obtained from CE2 and DCC, DESeq2 was used with a p-value<0.05 and |Log2FoldChange|>1. For visualisations of plots and graphs, R Studio packages were used. For generating overlapping circRNAs annotated from CE2 and DCC, the ‘Venn Diagram’ package was used. Histograms plotted to characterise the circRNA distribution across all chromosomes, putative circRNA count, length, start and end circularized exons were generated using ‘ggplot2’ from CE2 annotations. Circos plot for visualisation and frequency of circRNA expression was generated by ‘circlize’ package, and the heatmaps were generated using ‘pheatmap’. circRNA-miRNA interactions were observed using circBank database (https://www.circbank.cn/) and miRNA-mRNA interactions were studied using MultiMiR bioconductor package in R[45], [46].

### Cell culture

All cell cultures were performed at 37°C and 5% CO_2_ in humidified CO_2_ incubator under sterile environment.

### K562 cell culture, erythroid differentiation and HbF induction

Human erythroleukemia K562 cells were kindly gifted to us by Dr. Ramalingam from IGIB, New Delhi. Cells were maintained in RPMI 1640 (Gibco) supplemented with 10% Fetal Bovine Serum (FBS) (Biowest, VWR), 1% Glutamax (2mM Glutamine; Gibco) and 1% Penicillin and Streptomycin (Gibco). Erythroid differentiation of K562 cells was induced by 3U/ml EPO (PeproTech) for 72 hours. Fetal hemoglobin was induced by Hydroxuyrea (Sigma Aldrich, USA) at varied concentrations (50μM, 100μM, 200μM and 300μM) for 24 hours.

### *Ex vivo* culture of human CD34+ cells

The present study was approved by the Institutional Ethics Committee of Indian Institute of Technology, Kharagpur (ethics no. IIT/SRIC/DEAN/2024, date: 08.08.2024). Peripheral blood mononuclear cells (PBMC) were isolated by FicolPaque (Cytiva) density gradient centrifugation and the buffy coat containing the mononuclear cells were separated. CD34+ cell enrichment from mononuclear cells was performed by CD34 MicroBead Kit Ultrapure, human (#130-100-453; Miltenyi Biotec) according to manufacturer’s protocol. CD34+ hematopoietic stem and progenitor cells (HSPCs) were proliferated in StemSpan SFEM II medium supplemented with 100ng/ml SCF (PeproTech), 20ng/ml IL-3 (Miltenyi Biotec), 50ng/ml TPO (R&D Bioscience), 100ng/ml Flt3-Ligand (PeproTech), 2mM L-Glutamine (Gibco) and 1% Penicillin-Streptomycin (Gibco). To induce differentiation, *ex vivo* CD34+ HSPCs were cultured in basal medium consisting of Iscove’s Modified Dulbecco’s Medium (Gibco), 5% human AB serum (Gibco), 2mM L-Glutamine, 1% Penicillin-Streptomycin, 10µg/ml Insulin (Sigma), 330µg/ml Holo-Transferrin (R&D Bioscience), and 3U/ml Heparin (HiMedia). The medium was supplemented with 3U/ml EPO (PeproTech), 20ng/ml SCF (PeproTech), 1ng/ml IL-3 (PromoKine) and 100µM Hydroxyurea (HU) for induction of fetal hemoglobin.

### HEK293T cell culture

HEK293T cells were obtained from National Centre for Cell Science (NCCS), Pune. Cells were cultured in Dulbecco’s Modified Eagles Medium (DMEM; Gibco), supplemented with 10% FBS and 1% Penicillin and Streptomycin. Cells were detached for passaging by using 0.5% Trysin-EDTA (Gibco, USA) for 2 mins at 37°C.

### Fluorescence-activated cell sorting (FACS) analysis

For studying erythroid differentiation, cells were collected and washed with ice-cold phosphate-buffered saline (PBS) with 2% FBS. 1×10^5^ cells were stained with PE-conjugated anti-CD71 (#130-115-029; Miltenyi Biotec) and FITC-conjugated anti-CD235a antibody (#130-117-688; Miltenyi Biotec) in PBS/2% FBS at 4°C for 30 minutes. For studying the F-cell population, 2-3×10^5^ cells were fixed with 0.05%(v/v) glutaraldehyde for 10 min and permeabilized with 0.1% Triton X-100 (Sigma) for 5 min at room temperature. Cells were stained with APC-conjugated HbF antibody (#130-126-126; Miltenyi Biotec) on ice for 30 min. Cells for all assays were finally washed with PBS/2% FBS and subjected to Flow cytometric analysis on FACS LSRII flow cytometer (BD Biosciences). All data were analyzed by FlowJo v11.

### Morphology study by Leishman stain

For morphological analysis, primary erythroid cells were harvested and smeared on glass slides by cytocentrifugation (SLEE CS Centrifuge Type I) at 100 x g for 5 minutes. After air drying for 5 mins, slides were stained with Leishman stain (Sigma) solution for 5 minutes followed by incubation with double the amount of buffered water (pH 7.2) for 10 min. Slides were washed repeatedly with distilled water to remove excess stain and air dried. Images of the stained cells were taken using Nikon TE2000 microscope equipped with a digital camera.

### Plasmid construction, miRNA mimics and siRNA transfections

The pcDNA3.1(+) CircRNA Mini Vector was a gift from Jeremy Wilusz (Addgene plasmid #60648; http://n2t.net/addgene:60648; RRID: Addgene_60648)[47]. For overexpression of circNFATc3, the cDNA of NFATc3 exons 2 and 3 were amplified from K562 cells and cloned into pcDNA3.1(+) CircRNA Mini Vector and pcDNA3.1(+) empty vector (Supplementary Table S1). K562 cells were transfected with the overexpression vectors using Lipofectamine 2000 (Invitrogen, USA). Following transfection, G418 (Sigma) selection was performed for at least two weeks in a 96-well plate. Confirmation of overexpression was assessed by RT-qPCR. miScript miRNA mimics (hsa-let-7b-5p #YI04100945-ADA) and mimic negative control (miR-NC #YI00199006-ADA) were purchased from Qiagen, Germany. Custom-designed siRNA against the backsplice junction of circNFATc3 (si-circNFATc3) and siRNA negative control (si-NC) were obtained from Barcode Biosciences (Supplementary Table S2). miRNA mimics (final concentration of 100nM) or siRNA transfections (final concentrations of 25, 50, 100 and 150 pmols) were performed using Lipofectamine 2000. Cells were incubated for 8 hours in Opti-MEM and replaced with RPMI medium supplemented with 10% FBS and 2mM L-Glutamine, and incubated for 48 hours before proceeding with further studies.

### Real-time quantitative PCR

Total RNA was extracted using miRneasy Mini Kit (Qiagen, Germany) according to the manufacturer’s protocol. For qPCR of mRNA and circRNA, 1µg of total RNA was reverse transcribed using High-Capacity cDNA Reverse Transcription kit (Invitrogen, USA) with random hexamer primers. miRNA cDNA conversion was performed by converting 1µg of total RNA using miRCURY LNA RT II Kit (Qiagen, Germany). For qPCR analysis, PowerUp SYBR Green Master Mix (Applied Biosystems, USA) was used to detect mRNAs and circRNAs according to the manufacturer’s protocol. miRNA qPCR was performed by miScript primer assay (hsa-let-7b-5p #MS00003122; hsa-RNU6-2_11 #MS00033740; Qiagen, Germany) using stem-loop primers according to the manufacturer’s protocol. The expression of mRNAs and circRNAs was normalised by β-actin, and miRNA expression was normalised by U6 snRNA. All primers used in qPCR are enlisted in Supplementary Table S3. All circRNA-specific divergent primers were designed in Circinteractome (www.circinteractome.irp.nia.nih.gov), as per the guidelines.

### Immunoblot analysis

For immunoblot analysis, cells were lysed using RIPA buffer (Sigma), containing cOmplete Protease Inhibitor Cocktail (Roche), and the total protein was quantified by Pierce BCA Assay Kit (Invitrogen, USA). Protein samples were prepared using 2X Lammelli sample buffer (Bio-rad) and heated at 95°C for 10 minutes, following which the protein was separated based on their molecular weight by 12% SDS-PAGE. The separated proteins were transferred into a nitrocellulose membrane and blocked by 5% (w/v) skim milk. The membrane was incubated overnight at 4°C with specific primary antibodies, which included, anti-HBG antibody (#39386, CST), anti-GATA2 antibody (#4595, CST), anti-β-actin antibody (#ab8227, abcam) and anti-H3 antibody (#A2348, abclonal). The membranes were washed and incubated with secondary antibody goat anti-rabbit IgG H+L-HRP (#ab6721) for 2 hours at room temperature. The band were detected by Clarity Western ECL Substrate (Bio-rad) and ChemiDoc (Bio-rad). In this study, β-actin and Histone H3 were used as internal controls.

### Luciferase reporter assay

Single-stranded forward and reverse strand oligos having let-7b-binding sequences along with their flanking sequences in circNFATc3 and GATA2 3’UTR were obtained from Integrated DNA Technologies (IDT). All oligo sequences have been listed in Supplementary Table S4. The forward and reverse strands of the oligos were annealed in annealing buffer (10mM Tris HCl, pH 7.5-8.0, 50mM NaCl, 1mM EDTA) at gradient temperature. The oligos were designed to contain overhangs of XbaI site in the forward strand and SacI site on the reverse strand. The double-stranded oligos were cloned into the XbaI and SacI site of pmirGLO Dual-Luciferase miRNA Target Expression Vector (#E1330; Promega, USA), at the C-terminal of *luc2* cassette. The mutated oligos were designed by changing the let-7b—binding regions to create mutant vectors. The recombinant wild type or mutant plasmids were co-transfected with hsa-let-7b-5p mimics in HEK293T cells using Lipofectamine 2000 (Invitrogen, USA). After 24 hours of transfection, luciferase activity was assessed using Dual GLO Luciferase Reporter Assay System (Promega, USA) in Glomax Luminometer (Promega, USA) according to manufacturer’s protocol. The firefly luminescence signals were normalised against *hRluc* cassette signal.

### Validation of circular structure of RNA

To detect the presence of circNFATc3, total RNA was extracted from K562 cells using the miRNeasy Mini Kit (Qiagen, Germany). Subsequently, the RNA was treated with RNase R, an exonuclease capable of degrading linear RNAs while preserving circular structures[48]. Briefly, 1 U of RNase R (Abcam) was used to treat 1 µg of total RNA for 30 minutes at 37°C, followed by a heat inactivation at 65°C for 20 minutes. Reverse transcription was performed, and the abundance of circNFATc3, NFATc3, and β-actin in RNase R-treated and untreated RNA was quantified by RT-qPCR.

### Pull-down assay with Bio-let-7b-5p

Synthetic let-7b-5p mimics and mimic negative control (miR-NC) were Biotinylated at their 3’ ends using the Pierce RNA 3’ End Biotinylation Kit (Invitrogen, USA) according to manufacturer’s protocol. RNA-pull down was performed on K562 cells overexpressing circNFATc3 by the method described previously[49]. Briefly, cells were fixed with 1% paraformaldehyde (PFA) and lysed by sonication using lysis buffer (50 mM Tris-HCl pH 7.0, 1% SDS, 10 mM EDTA, 200 U/mL RNAse inhibitor, and 5 µL/mL protease inhibitor cocktail). 100 pmol of the biotinylated let-7b-5p mimics or miR-NC were added and incubated in hybridisation buffer (50 mM Tris-HCl pH 7.0, 1 mM EDTA, 750 mM NaCl, 15% Formamide, and 1% SDS). Dynabeads M-280 Streptavidin (∼1mg) (Invitrogen, USA) was added to the suspension to promote biotin-streptavidin interactions and bead recovery was done using magnetic support. RNA recovery was done using Proteinase K buffer (10 mM Tris-HCl pH 7.0, 100 mM NaCl, 1 mM EDTA, 0.5% SDS) and 5 µL of proteinase-K (20 mg/mL) and heated at 50°C for 45 mins and 95°C for 10 mins. RNA was isolated by the miRNeasy RNA isolation kit (Qiagen, Germany) and reverse transcribed. RT-qPCR was performed to evaluate the enrichment of circNFATc3.

### Sub-cellular fractionation study

K562 cells were washed with PBS and resuspended in ice-cold cytoplasmic extraction buffer [0.05% (v/v) NP-40, 0.5mM DTT, 1mM EDTA, 10mM HEPES (pH7.9), 10mM KCl and 1.5mM MgCl_2_] and incubated for 5 min on ice. Cell lysate was then centrifuged at 16,000 g for 10 min at 4 °C and supernatant was collected as cytoplasmic fraction. The pellet was resuspended with nuclei extraction buffer (26% (v/v) Glycerol, 0.2mM EDTA, 0.5mM DTT, HEPES (pH7.9), 300mM NaCl and 1.5mM MgCl_2_] and incubated for 5 min on ice. The lysate was centrifuged 25,000 g for 10 min. The supernatant was collected as nuclear fraction. RNA was extracted from both nuclear and cytoplasmic fractions using TRIzol (Applied Biosystems) method according to manufacturer’s protocol. RT-qPCR was performed to measure the abundance circNFATc3 in subcellular fractions. The expression of Malat1 and β-actin were regarded as internal control for nuclear RNA and cytoplasmic RNA, respectively.

### Cell viability assay

To evaluate the cell viability after all transfections, K562 cells were harvested from 12-well plate and seeded in 96-well opaque culture plate (Corning). CellTiter-Glo® reagent (Promega) was added according to manufacturer’s protocol to each well and the luminescence signal was read after 15 minutes with GloMax® Navigator Microplate Luminometer (Promega).

### Total RNA sequencing

Total RNA was isolated from K562 cells overexpressing circNFATc3 in pcDNA3.1(+) circRNA mini vector and NFATc3 (exons 2-3) linear isoform in pcDNA3.1(+) plasmid using miRNeasy RNA isolation kit (Qiagen, Germany). Paired-end total RNA sequencing was performed for 3 biological replicates for both control and treated samples in NovaSeqX Plus platform, with 150bp read length, and 100 million reads per sample (MedGenome Labs Pvt Ltd.).

After the raw reads were extracted, quality reports were used to trim reads. Quality control was performed by FastQC (v0.11.9), while adapter removal utilised fastq-mcf (v1.05) and cutadapt (v4.7). To prepare the RNA-Seq data, non-target sequences such as mitochondrial DNA, ribosomal RNAs, transfer RNAs, and adapter sequences were filtered out using Bowtie2 (v2.5.3). Paired-end sequencing reads were mapped against the human reference genome (GRCh38/hg38) using STAR aligner (v2.7.11b) with the relevant FASTA and GTF annotation files obtained from the Gencode database.

Gene expression was normalised in FPKM using cufflinks (v2.2.1). DESeq2 was employed for detecting differentially expressed genes. Heatmap and volcano plot were generated by ‘pheatmap’, ‘ggplot2’ and ‘ggrepel’. Gene ontology (GO) and pathway analysis was performed by Overrepresentation Analysis (ORA) and KEGG pathway analysis. ORA was applied with organism-specific gene annotation (OrgDb), filtering enriched categories by p-value (<0.05) and controlling for FDR. Enrichment scoring for KEGG and GO categories utilised the clusterProfiler methods enrichGO, enrichKEGG, gseGO, and gseKEGG, with up-to-date gene annotation and pathway databases and ‘ggplot2’ was used for visualisation of the dotplots and networks.

### Statistics and reproducibility

All data were analyzed using Graphpad prism v10.2 software and they have been represented as Mean ± SEM. Statistical differences were evaluated by using Student t*-*test. All experiments have been performed at least three times, and a p-value <0.05 was defined as statistically significant.

## ETHICAL APPROVAL

The ethical approval for the study was obtained from IIT Kharagpur prior to sample collection. (Approval No. IIT/SRIC/DEAN/2024, date: 08.08.2024).

## DATA SHARING STATEMENT

The data are not publicly available due to privacy or ethical restrictions. For original data, please contact-.

## AUTHOR CONTRIBUTIONS STATEMENT

M.M. and N.C. conceptualized the study. M.M. performed the in silico analyses. M.M. coordinated the blood sample collection, which was carried out by trained clinical personnel. M.M. and M.R. were involved in data collection, curation, and analysis. M.M. prepared the figure panels and wrote the manuscript. N.C. and T.K.D. reviewed and edited the manuscript. All authors have read and approved the final version of the manuscript.

## ACKNOWLEDGEMENTS

The authors would like to acknowledge School of Medical Science and Technology, Indian Institute of Technology Kharagpur for providing the support and infrastructure. The authors would also like to thank Dr. Sivaprakash Ramalingam from CSIR-Institute of Genomics and Integrative Biology (CSIR-IGIB) for providing the K562 cells. The authors would also like to acknowledge Dr. Gayatri Mukherjee’s lab at the School of Medical Science and Technology (SMST), IIT Kharagpur, for their invaluable assistance with flow cytometry. The authors would also like to thank Dr. Jeremy Wilusz for providing the pcDNA3.1(+) circRNA mini vector (Addgene plasmid #60648). The authors also acknowledge Perplexity AI, a large language model for its assistance in improving the clarity and language of the manuscript. Ms. Mandrita Mukherjee would like to acknowledge Ministry of Education, Government of India for financial support. This work has been supported by Anusandhan Nation Research Foundation (ANRF), Ministry of Science and Technology, Government of India. The project is titled as “Mechanistic insights on long non-coding RNA (lncRNA)-mediated ceRNA network in fetal hemoglobin regulation” (Sanction no.: CRG/2023/003509).

