## Supplementary data 8 for "CircNFATc3 promotes Fetal Hemoglobin induction by regulating let-7b/GATA2 axis"

**Table S1:** Primers used for amplifying 1298bp NFATc3 exon 2 and 3.

| Gene | Forward primer (5' — 3') [EcoRI overhang] | Reverse primer (5' — 3') [XhoI overhang] |
|---|---|---|
| **NFATc3** | GGAATTCATCTTGAGCCAGATGATTGTGC | CCGCTCGAGCTTCACAACAGGATGTCCC |

**Table S2:** Sequence of siRNA used against circNFATc3 BSJ.

| siRNA | Sequence (5' — 3') |
|---|---|
| **si-circNFATc3** | CCUGUUGUGAAGAUCUUGAGCUU |

**Table S3:** List of primers designed for RT-qPCR analysis.

| Gene | Forward Primer (5' — 3') | Reverse Primer (5' — 3') |
|---|---|---|
| hsa_circ_0000711 | TGAAACTGAAGGTAGCCGAGGG | TGGCTCAAGATCTTCACAACAGG |
| hsa_circ_0000568 | AGCTGCACCAGTGTTATTGGA | TGCTACCTTGATCTGAGCGT |
| hsa_circ_0075796 | TGGCCAGTGCTAACTTGGAAGAA | TCCCTCTTGCCTTCCACGATG |
| hsa_circ_0008032 | TGCTGGTAGCCTGTCAACAA | AGCACCAAATCCTCTACACAGT |
| γ(A+G)- globin | AAGCTCCTAGTCCAGACGCC | AGACAACCAGGAGCCTTCCC |
| GATA2 | CCAAGCGAAGACTGACGACAAC | TCTTCCGGTTCCGAGTCTGG |
| NFATc3 | TACCCGTTGAGTGCTCCCAG | CCCACTGAGGTCGTCCATCT |
| β-actin | ACTGGAACGGTGAAGGTGACA | AGTCCTCGGCCACATTGTGAA |
| MALAT1 | AGCAAACTGTGTTGGCGTGG | CGGTGCCTTTAGTGAGGGGT |
| GATA1 | GCCACTACCTATGCAACGCC | CCCGTTTACTGACAATCAGGC |
| ALAS2 | AGGAAGCCATTTTCCGGTCC | ACTGAAGACATAGTTTCCAGGC |
| BAND3 | AACGTAGCTGGTCGCAGAG | TGTCTACGGTGATCTGAGCC |
| HBB | TGGATGAAGTTGGTGGTGAG | CCTTAGGGTTGCCCATAACA |

**Table S4:** Sequences of oligos used for luciferase reporter assay.

| Gene | Forward oligo (5' — 3') [XbaI overhang] | Reverse oligo (5' —3') [SacI overhang] |
|---|---|---|
| **circNFATc3** | CGCGGCCGCTATATTTCGCACATCTTCATTACCTCCACTAGACTT | CTAGAAGTCTAGTGGAGGTAATGAAGATGTGCGAAATATAGCGGCCGCGAGCT |
| **NC-circNFATc3** | CGCGGCCGCTATATTTCGCACATCTTCATTAATTCCACTAGACTT | CTAGAAGTCTAGTGGAATTAATGAAGATGTGCGAAATATAGCGGCCGCGAGCT |
| **GATA2** | CGCGGCCGCGCAGATTTGTGGGGACCTCAGCCTGCACT | CTAGAGTGCAGGCTGAGGTCCCCACAAATCTGCGCGGCCGCGAGCT |
| **NC-GATA2** | CGCGGCCGCGCAGATTTGTGGGGATATCAGCCTGCACT | CTAGAGTGCAGGCTGATATCCCCACAAATCTGCGCGGCCGCGAGCT |
