## Supplementary data 9 for "CircNFATc3 promotes Fetal Hemoglobin induction by regulating let-7b/GATA2 axis"

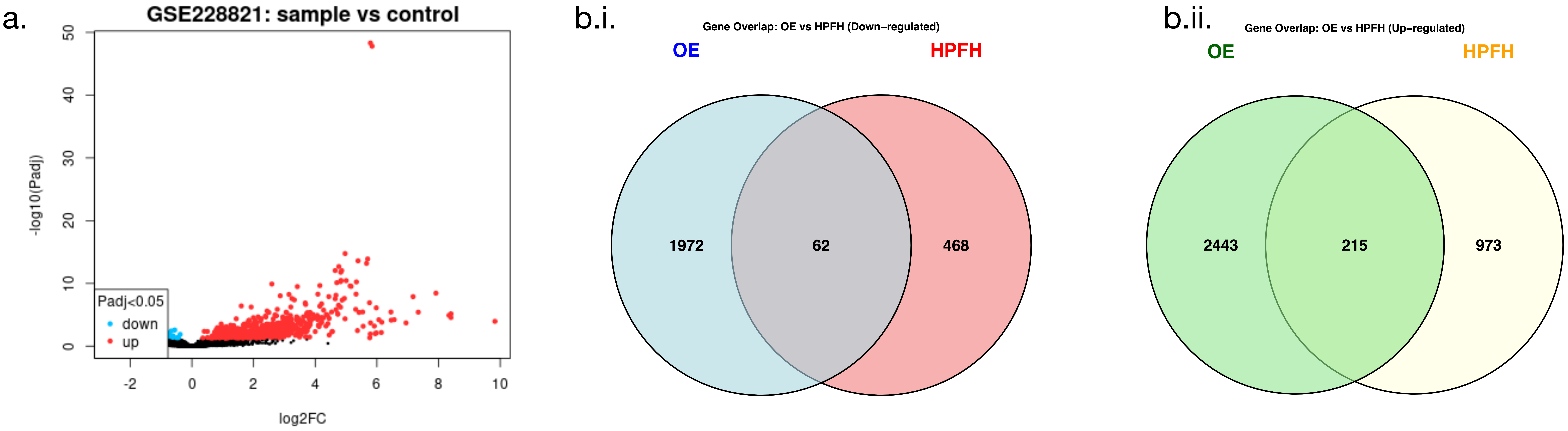


**Figure 1: a.** Volcano plot generated from dataset GSE228821 describing HSPCs have been base edited using CRISPR to mimic HPFH mutation and HbF induction. **b.** Overlapping genes that are **i.** downregulated and **ii.** Upregulated between dataset GSE228821 (HPFH) and circNFATc3 OE dataset obtained from our study.

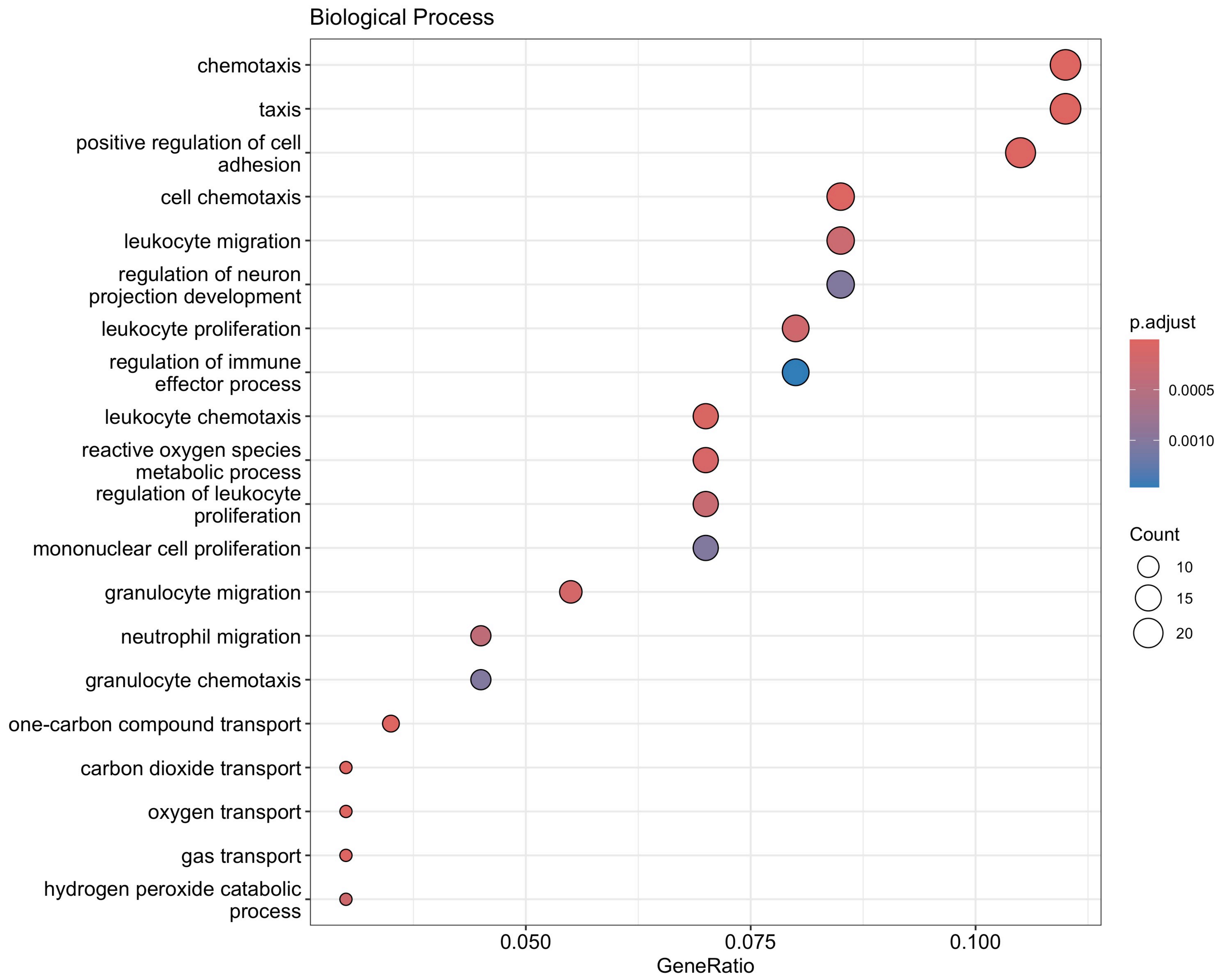


**Figure 2:** Biological process (BP) enriched by Gene Ontology (GO) study of overlapping genes between dataset GSE228821 (HPFH) and circNFATc3 OE dataset obtained from our study.

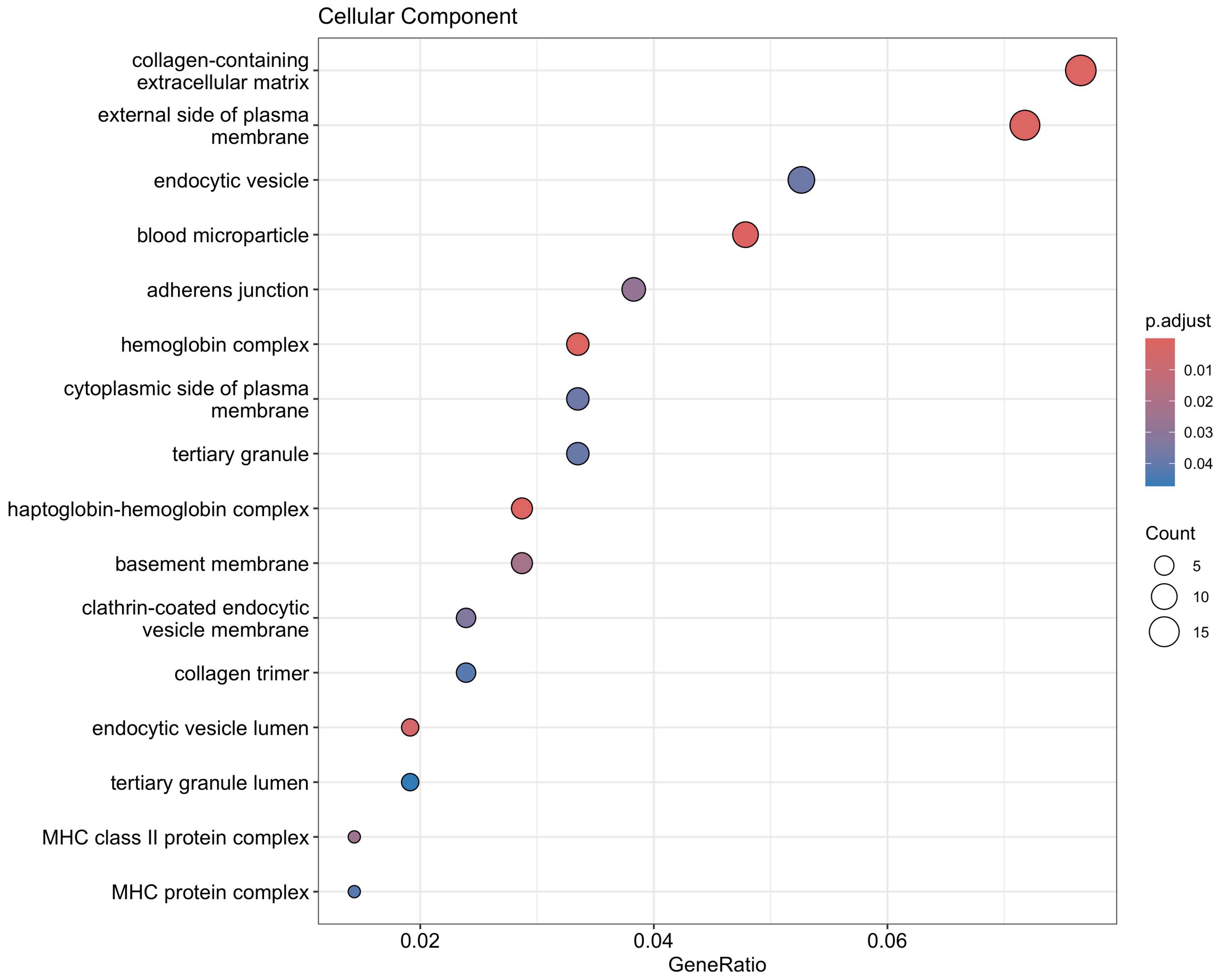


**Figure 3:** Cellular component (CC) enriched by Gene Ontology (GO) study of overlapping genes between dataset GSE228821 (HPFH) and circNFATc3 OE dataset obtained from our study.

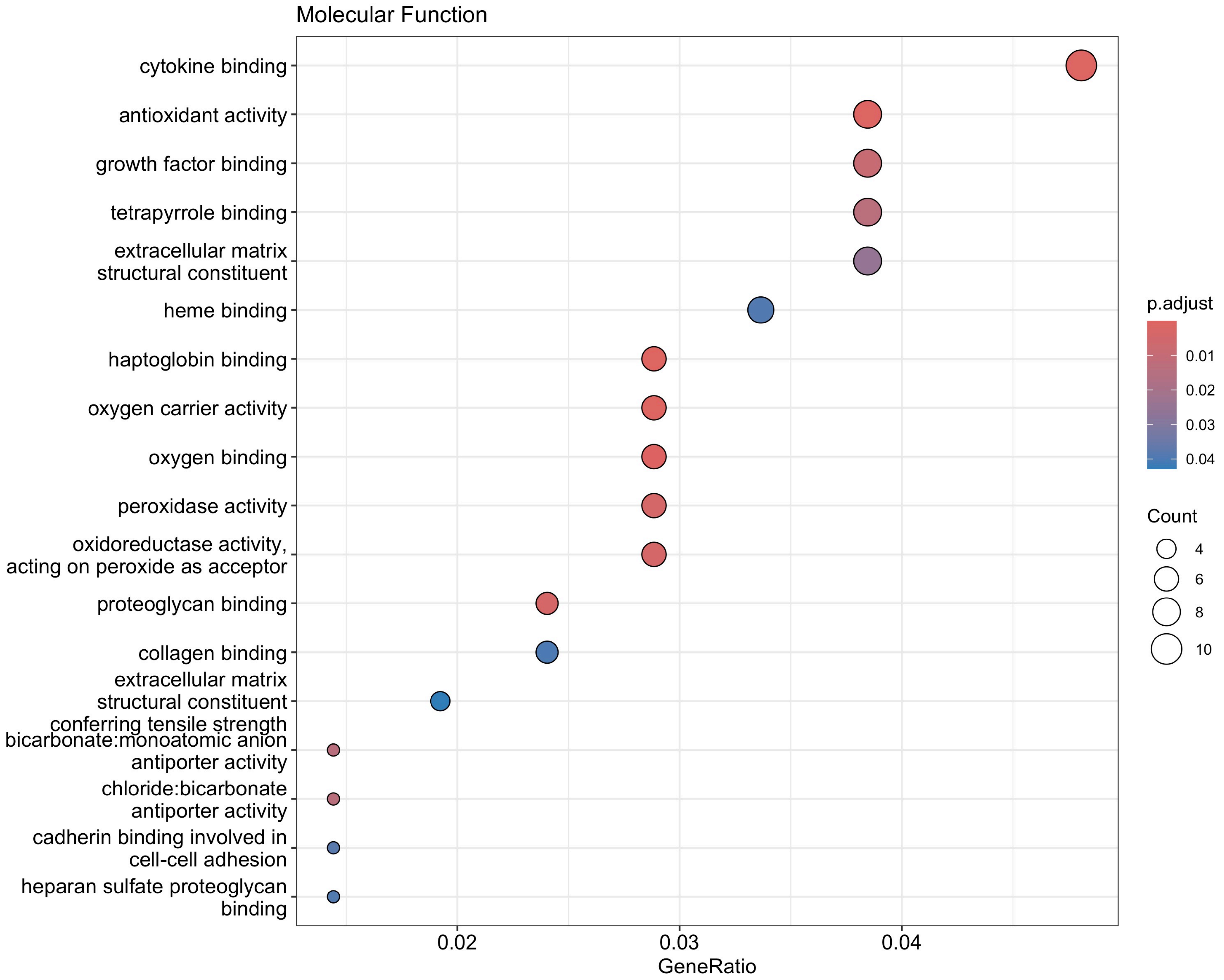


**Figure 4:** Molecular Function (MF) enriched by Gene Ontology (GO) study of overlapping genes between dataset GSE228821 (HPFH) and circNFATc3 OE dataset obtained from our study.

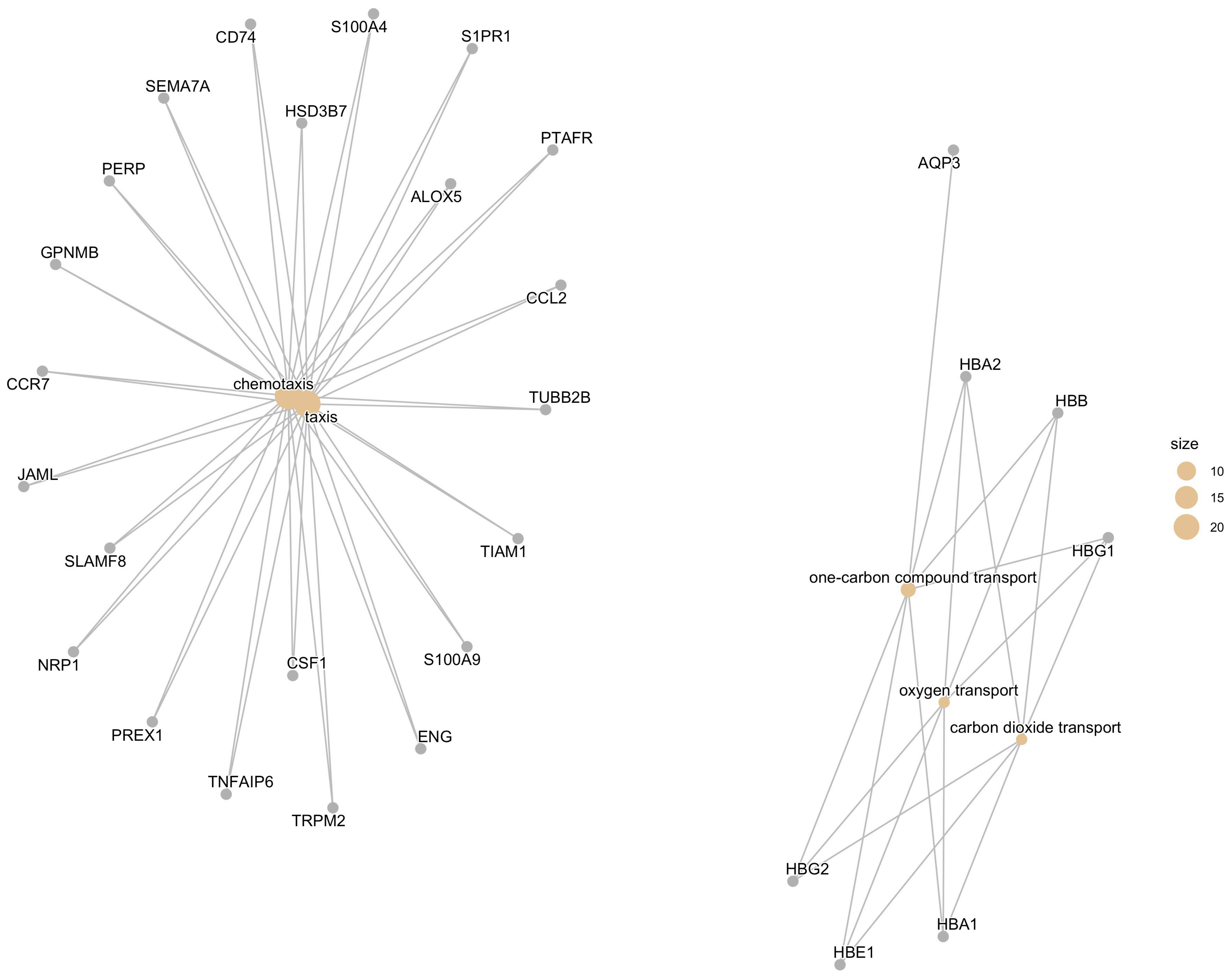


**Figure 5:** Gene-concept network of overlapping genes between dataset GSE228821 (HPFH) and circNFATc3 OE dataset obtained from our study.

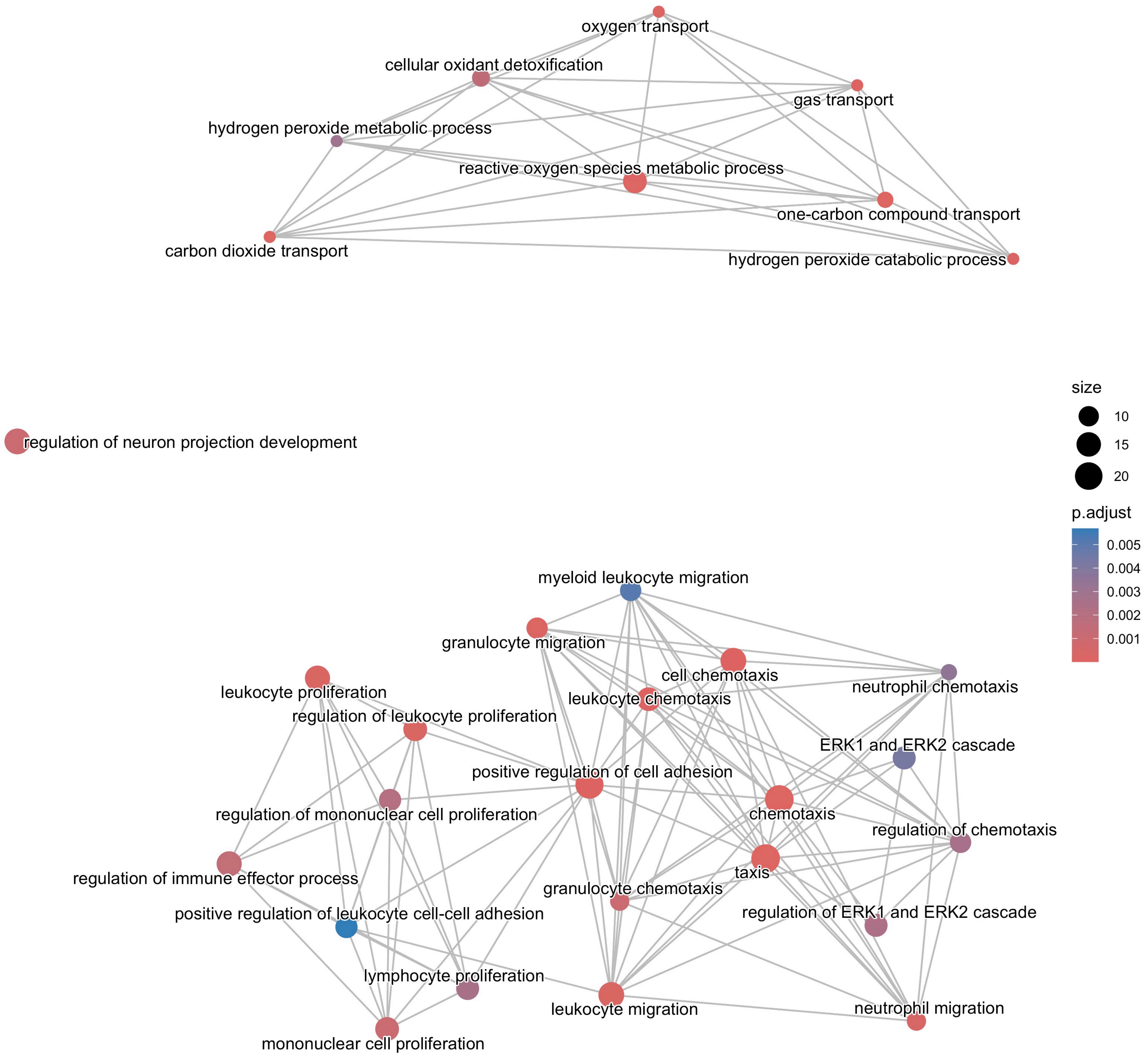


**Figure 6:** Enrichment map of overlapping genes between dataset GSE228821 (HPFH) and circNFATc3 OE dataset obtained from our study.
